# Drug resistance through ribosome splitting and rRNA disordering in mycobacteria

**DOI:** 10.1101/2024.06.13.598844

**Authors:** Soneya Majumdar, Amuliya Kashyap, Ravi K. Koripella, Manjuli R. Sharma, Kelley Hurst-Hess, Swati R. Manjari, Nilesh K. Banavali, Pallavi Ghosh, Rajendra K. Agrawal

## Abstract

HflX is known to rescue stalled ribosomes and is implicated in antibiotic resistance in several bacteria. Here we present several high-resolution cryo-EM structures of mycobacterial HflX in complex with the ribosome and its 50S subunit, with and without antibiotics. These structures reveal a distinct mechanism for HflX-mediated ribosome splitting and antibiotic resistance in mycobacteria. In addition to dissociating ribosome into two subunits, mycobacterial HflX mediates persistent disordering of multiple 23S rRNA helices to generate an inactive pool of 50S subunits. Mycobacterial HflX also acts as an anti-association factor by binding to pre-dissociated 50S subunits. A mycobacteria-specific insertion in HflX reaches further into the peptidyl transferase center. The position of this insertion overlaps with ribosome-bound macrolides or lincosamide class of antibiotics. The extended conformation of insertion seen in the absence of these antibiotics retracts and adjusts around the bound antibiotics instead of physically displacing them. It therefore likely imparts antibiotic resistance by sequestration of the antibiotic-bound inactive 50S subunits.

## Introduction

Mycobacteria are a complex group of bacteria with serious implications for human health. Their ability to cause diverse diseases and rapidly develop resistance to antibiotics makes treatment very challenging. Therefore, there is an increasing need to investigate the mechanisms of drug resistance in mycobacteria to develop improved treatment strategies and therapies. Previous studies^1^ have shown that the macrolide-lincosamide resistance in *Mycobacterium abscessus* (*Mab)* and *Mycobacterium smegmatis* (*Msm*) is associated with the High Frequency of Lysogenization X (HflX) protein. HflX is a universally conserved GTPase belonging to the Obg-HflX superfamily, and it resembles bacterial ribosome-binding GTPases involved in ribosome biogenesis ^2^. Its expression is induced under specific stress conditions such as heat, oxidative stress, or antibiotic treatment ^1, 3–6^. HflX was thought to be just a ribosome biogenesis factor ^2, 7^ until the *Escherichia coli* (*Eco*) HflX was shown to split 70S ribosomes into their large 50S and small 30S ribosomal subunits (henceforth refer to as LSU and SSU, respectively) during heat stress ^3^. HflX in *Staphylococcus aureus* (*Sau*) even recycles hibernating 100S disomes (dimers of 70S ribosomes) into the component LSU and SSUs ^8^.

Certain bacteria, like *Listeria monocytogenes* (*Lmo*), possess a second HflX gene, HflXr, which is implicated in resistance to macrolide, lincosamide, pleuromutilin, and streptogramin antibiotics ^5, 9^. While HflXr is suggested to confer resistance by allosterically modifying the drug target site on the ribosome ^5^, the single HflX protein in *Eco* is proposed to be involved in both stalled ribosome splitting and antibiotic displacement from the peptidyl transferase center (PTC) ^3, 6^. Like *Eco* and unlike *Lmo*, mycobacteria have a single HflX protein ^1^. In slow-growing *Mycobacterium bovis* (*Mbo*), HflX regulates the transition to a non-replicating, drug-tolerant state under hypoxia ^4^. In fast-growing species like *Mab* and *Msm*, unlike both *Eco*-HflX and *Lmo*-HflXr, HflX mediates resistance against macrolide-lincosamide antibiotics by splitting and recycling stalled ribosomes ^1^. Although regulation of HflX expression is similar across bacterial species, the proposed mechanisms of HflX-mediated antibiotic resistance vary.

Similar to most HflX homologs, mycobacterial HflX exhibits a three-domain architecture consisting of an N-terminal domain (NTD), a GTPase domain (GD), and a C-terminal domain (CTD) ^2, 3^. The NTD is composed of two subdomains: ND1, which interacts with the 23S rRNA helix 69 (H69, henceforth the 23S and 16S rRNA helices are identified by a number prefixed with H or h, respectively), and a fork-like α-helical domain (HD) with a loop that extends toward the ribosomal PTC ^3^, that is thought to be involved in antibiotic resistance ^5, 6^. Mycobacterial HflX possesses a species-specific NTE (39 amino acid in *Msm*) and a variable-length insertion (9 amino acids in *Msm*) in the HflX-HD ^1^. Time-resolved cryogenic electron microscopy (TRCEM) of HflX-mediated ribosome splitting in *Eco* ^10^ and *Lmo* ^11^ suggests that rotation of SSU enabled by HflX leads to inter-subunit bridge disruption and 70S splitting. Mycobacterial ribosomes have two additional inter-subunit bridges ^12–15^ formed by SSU proteins bS6 and bS22 interaction with H54a and H70, respectively, which may influence their splitting mechanism.

Here we present cryo-EM structures of *Msm* 70S-HflX and 50S-HflX complexes that represent multiple intermediate states for HflX interaction with the ribosome. We captured four intermediates with a bound HflX from *Msm* 70S-HflX-GMPPCP complexes: a 70S- HflX complex, and three different 50S-HflX complexes (referred to as 50S-HflX-A, 50S- HflX-B, and 50S-HflX-C). We also determined HflX-bound ribosome structures in the presence of two antibiotics, erythromycin (Ery) and Chloramphenicol (Clm). These structures show the following differences for *Msm*-HflX-mediated ribosome splitting and antibiotic resistance from those reported for *Eco* and *Lmo*: (i) the initial binding conformation for HflX on the 70S ribosome, (ii) a large internal conformational change within the HflX HD, (iii) a functional role for the 39-aa HflX NTE in step-wise disordering of multiple helices within domain IV of the 23S rRNA, and (iv) conformational changes in HflX-HD to accommodate PTC-binding drugs. These collectively indicate that HflX-mediated splitting and macrolide-lincosamide resistance in mycobacteria have diverged significantly from previously postulated mechanisms.

## Results and Discussion

### Structures of the Mycobacterial 70S-HflX and 50S-HflX-A Complexes

In the presence of a 10-fold molar excess of *Msm* HflX-GMPPCP, approximately 90 % of particles comprised 70S ribosomes with a bound E-site tRNA (**Figure 1a, b**; fig. S1) while 5.4% represents LSUs bound to HflX (**Figure 1c, d**; fig. S1). A minor fraction (4.6 % of 70S ribosome particle images) carried a 70S-bound HflX molecule and was reconstructed to a resolution of 3.2 Å (**Figure 1a, b**; fig. S1-2a). Despite the partial disorder of HflX, especially in the ND1 domain (**Figure 1b**), a portion of NTD with a partial density for the HD was present along with distinct densities for the GD and CTD domains. This structure likely represents a complex showing an early encounter of mycobacterial HflX with the 70S ribosome.

**Figure 1.**
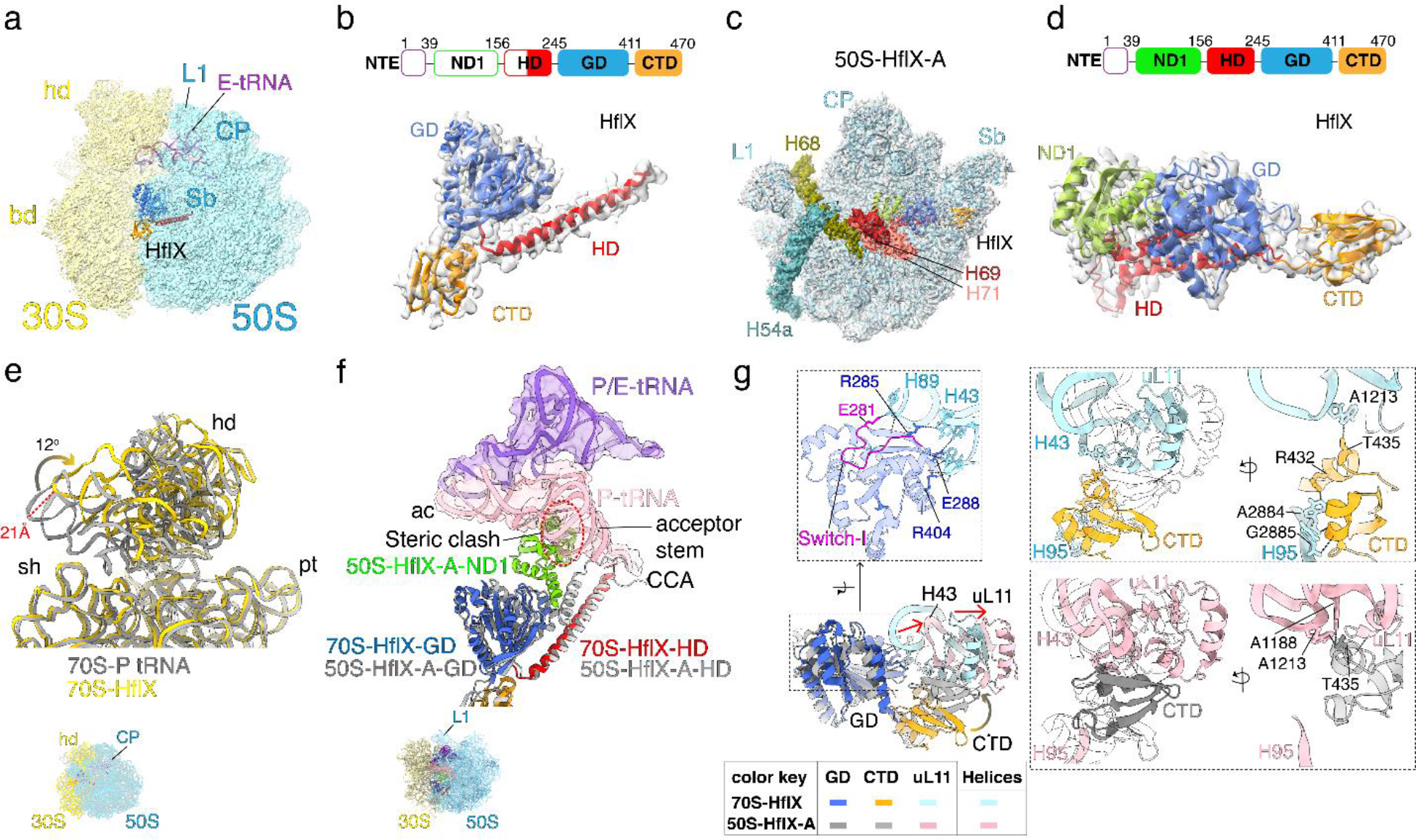
**Structures of the *M. smegmatis* 70S and 50S ribosome-HflX complexes.** (**a**) A 3.26 Å cryo-EM structure of the 70S-HflX complex showing well-defined HflX densities for its CTD (orange) and GD (blue) domains, and a partial density for its HD (red) domain, along with density for a deacylated E-state tRNA (purple). Structural landmark for 30S SSU (yellow): hd, head; bd, body; and for 50S LSU (cyan): L1, protein L1-stalk; CP, central protuberance, and Sb, L7/L12 stalk base. (**b**) Fitting of the molecular models of HflX domains into corresponding cryo-EM density. The top bar diagram depicts the domain architecture of *Msm* HflX, where solid color boxes indicate portions with visible HflX densities. Only half of the HD, corresponding to one of its two α-helices, was resolved in this structure. This color-coding convention is followed for all subsequent figures. (**c**) A 3.1 Å cryo-EM structure of the *Msm* 50S-HflX-A complex. Landmarks of the LSU: L1, protein L1 stalk; rest of the landmarks are as in panel a. The 23S rRNA helices H68 (olive green), H69 (brown), H71 (light salmon) and H54a (teal) are highlighted. **(d)** Modeling of HflX domains into their corresponding cryo-EM density. All domains of HflX, except the NTE, are resolved. **(e)** Superimposition of the 70S-HflX structure with the unrotated 70S-P tRNA structure. Shown are the 16S rRNA corresponding to head (hd) and portions of the body (shoulder, sh) and platform (pt) of the 70S-P tRNA (grey) and 70S-HflX (gold) structures. A 12° rotation of the SSU head in 70S-HflX complex as compared to that in 70S-P tRNA complex results in its lateral displacement by ∼21 Å. **(f)** Superimposition of the 70S-HflX structure with the structures of the 50S-HflX-A, *Eco* 70S bound to a P/E tRNA (PDB:7SSO), and 70S P- tRNA shows that the HflX-HD would extend to the CCA-end of the P-tRNA (light pink) while HflX- ND1 (green) domain would clash with the D-loop and acceptor arm of the P-tRNA, suggesting that full accommodation of the HflX-ND1 would require the P-tRNA to adopt the P/E state (purple). (**g**) Molecular interactions between the HflX-GD with H89 and H43. (i)The ordered Switch-I (magenta), a characteristic of the GTP bound ‘on’ state, seems to be monitored through its interaction with functionally relevant 23S rRNA helices. (ii) Superimposition of HflX from the 70S-HflX and 50S-HflX-A structures reveals that the HflX-CTD (orange) in the 70S-HflX structure interacts with the tip of H95 (α-sarcin-ricin loop, SRL) through its R432 residue, and (iii) subsequently shifts to the L7/L12 stalk base, as seen in the 50S-HflX-A complex to interact with H43 and protein uL11.

Structural comparisons between the 70S-HflX complex and the P-site tRNA bound 70S ribosomal class (fig. S1; **Figure 1e, f**) reveal conformational changes essential for initial accommodation of HflX on the ribosome. In the 70S-HflX complex, the SSU head undergoes a ∼12° rotation, moving the head towards the SSU platform by about 21 Å (**Figure 1e**) with concomitant transition of P-tRNA to P/E state (**Figure 1f**), apparently to accommodate HflX-ND1 into the 70S ribosome. If this rotation did not occur, the inner bend of the acceptor arm of any pre-bound P-site tRNA would sterically overlap with the HflX-ND1 domain (see **Figure 1f**).

A 3D reconstruction of LSU bound HflX, resolved to 3.1 Å, exhibited clear densities for all canonical HflX domains, except for the mycobacteria specific NTE (**Figure 1c, d**, fig. S1c, e, 2b). The loop connecting the two α-helices of the HD in *Msm* HflX is longer by 6 and 9-aa than the previously studied HflX orthologs in *Eco* and *Lmo*, respectively, and is well resolved in this structure (see fig. S3a, b). The overall conformation of *Msm* HflX and its binding to the LSU resembles the previously reported structures of *Eco* and *Lmo* HflX with their pre-dissociated LSUs ^3, 5, 6^.

**Figure 2.**
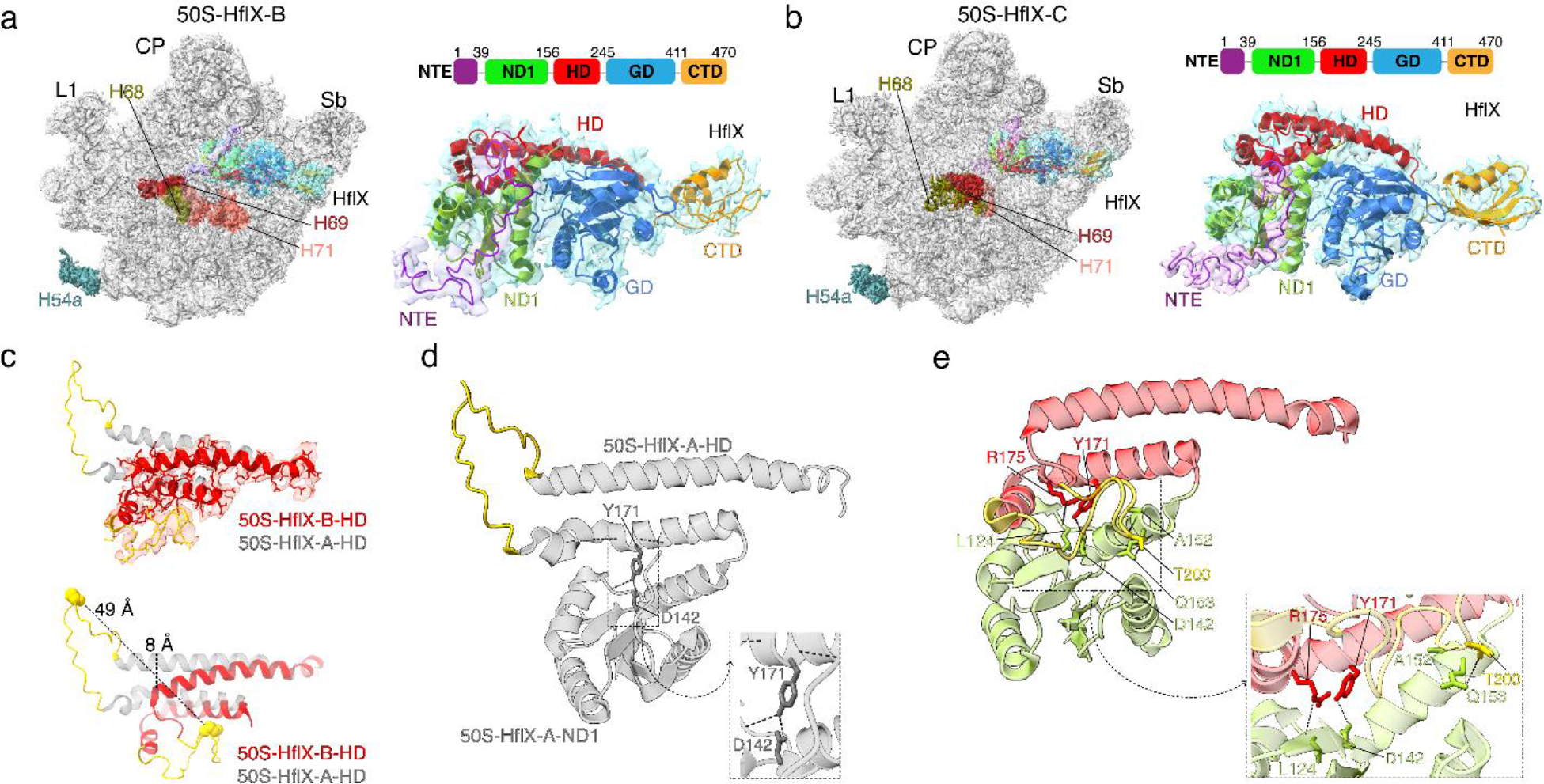
***Msm* HflX-binding induced splitting of the 70S ribosome is associated with a large conformational change in the protein** (**a**) Left- A 3.09 Å cryo-EM structure of *Msm* 50S-HflX-B complex. Landmarks of LSU are same as in Fig. 1c. The residual 23S rRNA helices H68-69, H71, H54a and the HflX domains are color-coded as previously. H68-69 and H54a are mostly disordered in this structure. Right-The fully resolved complete structure of *Msm* HflX bound to the LSU. (**b**) Left- A 2.96 Å cryo-EM structure of *Msm* 50S-HflX-C complex. H68-71 and H54a are mostly disordered in this structure. Right-The fully resolved complete structure of *Msm* HflX bound to LSU. The NTE in both 50S-HflX-B and 50S-HflX-C bar diagrams is shown in solid colors to reflect the fact that the NTE is also visible in both these states. (**c**) Superimposition of the HD from 50S-HflX-A (gray) and 50S-HflX-B (red) reveals two distinct conformations of the HD. The cryo-EM density corresponding to the bent conformation of the helical domain in 50S-HflX-B is displayed (top). The HD-loop is displaced by 49Å while the alpha-helix preceding the loop is moved by 8Å (bottom) (movie S1). (**d**) In the extended conformation (50S-HflX-A), interactions between HD and ND1 are feeble - Y171 of HD interacts with D142 of ND1 (inset). (**e**) In its retracted (folded) conformation (50S-HflX-B and 50S-HflX-C) interactions between the HD and ND1 are significantly increased- W179, R175 of HD interacts with the backbone of ND1-L124, Y171 of HD interacts with D142 of ND1 and, T200, V198 of HD interacts with Q153 of ND1. Insets show magnified views of the interactions.

A comparison between the 70S-HflX and the 50S-HflX-A complex shows a distinctly different positioning of the HflX-CTD on the ribosome. In the 70S-HflX complex, the CTD interacts with H95, the α-sarcin/ricin stem-loop (SRL). In the 50S-HflX-A complex, it interacts with protein uL11 and H43-H44 at the ribosomal stalk-base region (Sb) (**Figure 1g**; fig. S3c). In the 70S-HflX complex, the Sb itself moves (by ∼ 11 Å) and interacts with HflX-GD (**Figure 1g**). In both structures, a well-defined density for the GD Switch I-loop (**Figure 1g**; fig. S3d) and GMPPCP (fig. S3e) clearly indicates a pre-hydrolysis ’on’ state for HflX-GD, with the ordered Switch-I interacting with H89 (**Figure 1g**; fig. S3e).

HflX-HD interacts with H71 and H89-93 in the 70S-HflX complex (fig. S4a). In the 50S- HflX-A complex, HflX-HD interacts with H74, H89-H91, and the peptidyl transferase center (PTC) (fig. S4b), and ND1 is well resolved and lies adjacent to H69 (fig. S4c). As found earlier, the H69 tip moves upon HflX binding by ∼17 Å to cause a steric clash with the 16S rRNA h44 in the 70S ribosome structure (fig. S4d), thus inhibiting LSU’s association with SSU.

Above results suggest that *Msm*-HflX is recruited to the 70S ribosome through initial accommodation of its HD, GD, and CTD (**Figure 1a, b**), where HD extends to the CCA end of the P-tRNA likely to assess the acylation state of the tRNA (**Figure 1f**). The ribosomal Sb, known to be a dynamic structure that interacts with multiple translational factors ^16–22^, engages with the GD (**Figure 1g**). The GD-Switch I interacts with H89 (**Figure 1g**), potentially to recognize the GTP-bound state of the protein—an essential step for ribosome binding of HflX ^1, 3, 5, 10, 11^. SSU head rotation along with tRNA transition to P/E state facilitate HflX-ND1 binding, with concomitant interaction of HflX-CTD with the Sb (**Figure 1e-g**). The 50S-HflX-A complex represents a complete accommodated state of HflX onto the ribosome, where its ND1-mediated displacement of H69 in the LSU prevents ribosomal subunit association to form 70S ribosomes.

### Structures of the Mycobacterial 50S-HflX-B and 50S-HflX-C complexes

In the *Msm* 70S-HflX-GMPPCP complex prepared with a 20-fold molar excess of HflX, we observed significant (> 85 %) splitting of the 70S ribosomes. 3D variability analysis identified two distinct 50S-HflX states, referred to as 50S-HflX-B and 50S-HflX-C (refined to 3.09 Å and 2.96 Å resolution, respectively) (fig. S5). All domains of HflX were well resolved in these structures, however, the density corresponding to the 39-aa NTE required gaussian filtering to enable complete modeling (**Figure 2a, b**). A comparison to the 50S-HflX-A complex reveals the following differences with both these complexes: (i) density corresponding to the NTE of HflX is present; (ii) a large movement (∼49 Å) of the apical half of HD away from the PTC and onto the ND1 gives the HD domain a unique retracted and bent conformation (**Figure 2c-e**, movie S1) ; (iii) the H54a, H68 and H69 are disordered, with additional disordering of H71 in the 50S-HflX-C complex (**Figure 3**, movie S2); and (iv) HflX is deflected outward by ∼3.4 degrees, moving GD, HD and ND1 away from LSU. The pivot point for this deflection is on an axis along the CTD, which stays at the same location in all three 50S-HflX complexes (fig. S6a).

**Figure 3.**
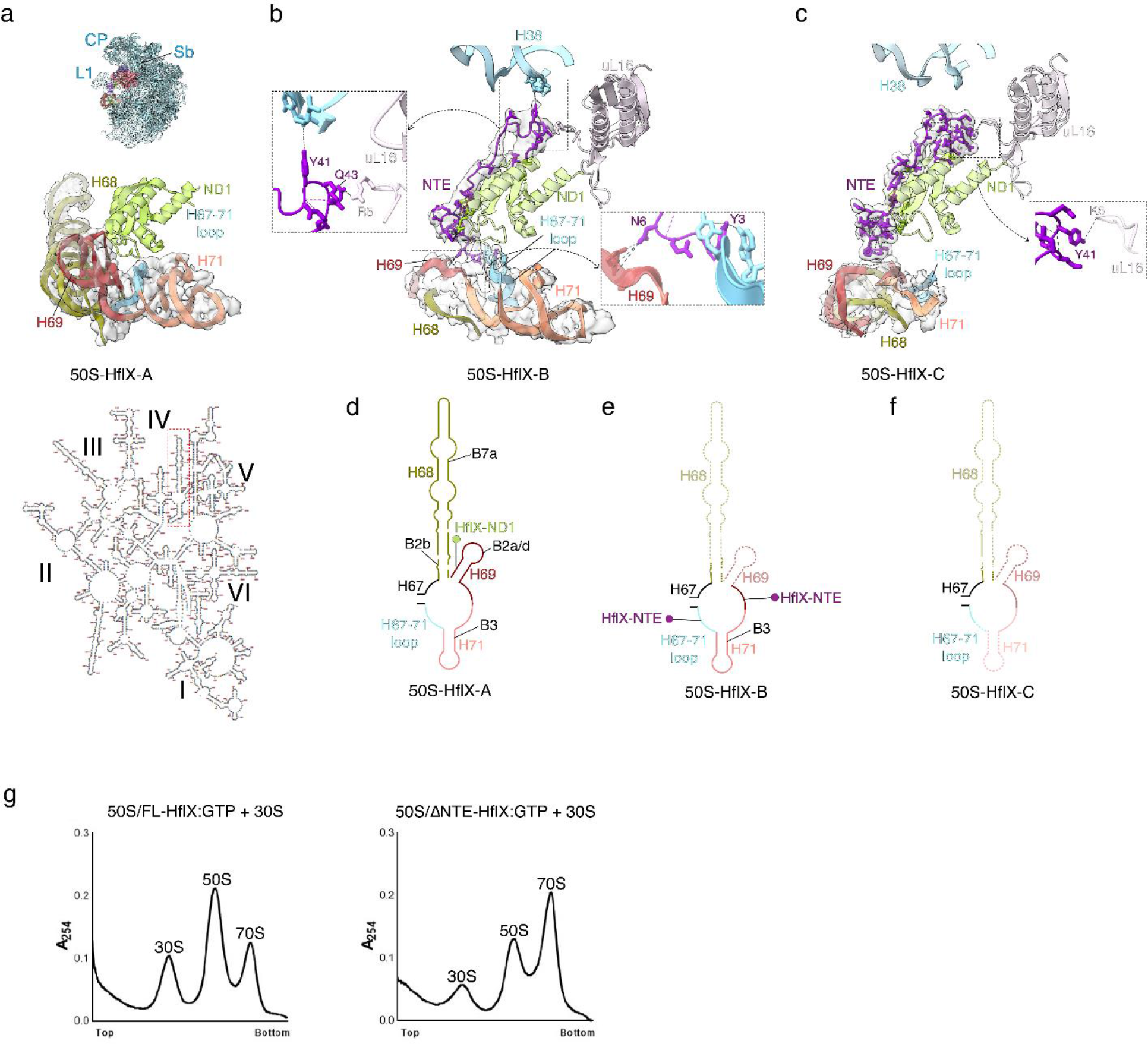
*Msm* HflX-binding induced splitting of the 70S ribosome is accompanied with sequential disordering of the 23S rRNA helices H68-71. (**a**) In 50S-HflX-A, H68 (olive green), H69 (brown) and H71 (light salmon) are in a well-resolved native conformation. **(b)** In 50S-HflX-B, HflX-NTE is stabilized through interactions with H38 (with Y41) and uL16 (with Q43) (inset). The N-terminal amino acid residues, N6 and Y3 of NTE interacts with H69 and the loop connecting H67 and H71, respectively. Concomitantly, H68 and H69 in this structure are disordered while H71 remains ordered. In 50S-HflX-C, the HflX-NTE loses interaction with H38 but the same aa residue (Y41) interacts with uL16 (inset). **(c)** While the interactions of the N-terminus of HflX-NTE seen in 50S-HflX-B are altered in 50S-HflX-C (also see fig. S6c), the entire H68-71 segment of the 23S rRNA is disordered (movie S2). **(d,e,f)** Schematic representation of the secondary structure of rRNA depicting the sequential disordering of helices across 50S-HflX states A, B and C along with the concurrent disruption of inter-subunit bridges. **(g)** Sucrose-gradient profile showing reassociation of *Mab* 50S subunits generated either with FL-*Mab* HflX (left) or *Mab* ΔNTE-HflX (right) in the presence of GTP with *Mab* 30S subunits. *Msm* and *Mab* HflXs show strong sequence homology (fig. S13) and are functionally exchangeable^1^.

The bending of HD’s α-helix in 50S-HflX-B and 50S-HflX-C occurs at a proline residue, P174, and a following stretch of glycines and alanines also likely enable the “retracted- bent” HflX conformation through increased flexibility. HD’s α-helix 2 mostly retains its interactions with the H89 and H90 (fig. S6b) but undergoes uncoiling at its terminal end from T206 to R213 and is deflected by 8 Å (**Figure 2c**). In 50S-HflX-A, the α-helix 2-Y171 residue interacts with the ND1-D142 residue. In 50S-HflX-B and 50S-HflX-C, the bent conformation of α-helix 2 makes multiple additional contacts with ND1 through the R175- L124 backbone, the T200-A152 backbone, and T200-Q153 (**Figure 2d, e**).

The NTE, unresolved in 50S-HflX-A, is stabilized in 50S-HflX-B through interactions of its Y41 residue with the A-site finger (U1013 of H38) and its Q43 residue with uL16 (R5) (**Figure 3b**). The rest of the NTE runs along the length of ND1, interacting with its ß2- strand and the adjacent loops, to extend towards the disordered junction of H68-69 (**Figure 3b, c;** fig. S6c). The NTE-Y3 residue interacts with the loop connecting H67 and H71 (residues A2190 and C2189), while the NTE-N6 residue interacts with the loop between H69 and H71 (the sugar-phosphate backbone of U2155-A2156, **Figure 3b**). In 50S-HflX-C, the NTE is arranged differently than in 50S-HflX-B. The NTE-Y41 residue interacts with uL16-R5 backbone but not the A-site finger (**Figure 3c**). The NTE extends along ND1 and interacts with its β2-α2 region and the loop connecting the two (fig. S6c). NTE interaction with the rRNA in 50S-HflX-C is uncertain since the H68-H71 segment is completely disordered in this structure (**Figure 3c, f**). The HflX-NTE is thus stabilized on the ribosome through A-site finger and uL16 interactions with concomitant bending of HflX-HD and retraction of HflX-GD and HflX-ND1 from the LSU surface.

A distinguishing feature is the disordering of four 23S rRNA helices in the 50S-HflX-B and 50S-HflX-C complexes. H54a, H68, and H69 are disordered in 50S-HflX-B, with H71 being additionally disordered in 50S-HflX-C (**Figure 3a-f**). These four 23S RNA helices form inter-subunit bridges B2a/d, B2b, B7a and B3 in the 70S ribosome ^12, 14, 15, 22–25^. The disorder in H71 in 50S-HflX-C, occurring in addition to disorder in H68 and H69, implies that these helices are disordered in a stepwise manner upon HflX binding (**Figure 3a-f**). We also observe a significant population of LSU without HflX, but with disordered H68- H71 (fig. S5). This suggests that these rRNA helices are disordered even after HflX dissociates from the ribosome, thus leaving behind an inactive pool of LSUs, resembling a late LSU biogenesis intermediate or an inactive LSU ^26–28^.

GTP hydrolysis has been suggested to be required for HflX release from the ribosome, rather than its splitting activity in eubacteria ^1, 3, 5^. Since our complexes were prepared with a non-hydrolyzable GTP analog GMPPCP, the observation of a significant population of HflX-free inactive LSUs (64 %) (fig. S5) suggests that the retraction of HflX-HD from the PTC (seen in 50S-HflX-B and 50S-HflX-C, fig. S6b), and NTE interaction with the disordered helices H68-69 could help deflect mycobacterial HflX away from the ribosome (fig. S6a), leading to release of HflX from LSU.

### Structure of the 50S-ΔNTE-HflX Complex

To confirm if HflX-NTE has a role in disordering of H68-71, we generated a ΔNTE-HflX construct that competently splits the 70S ribosomes in the presence of GMPPCP (fig. S7). The resulting 50S-ΔNTE- HflXcomplex, solved at a resolution of 3.4 Å, resembles the 50S-HflX-A complex, with the HD extended to the PTC and well-resolved H68-71 (fig. S7), confirming that NTE has a role in enabling disorder in H68-71 23S RNA helices. We hypothesized that the LSU pool generated by ΔNTE-HflX would be proficient in reassociation with SSU whereas that generated using full length HflX would be less proficient. To test this, we obtained post-dissociated LSUs that were split either with full-length-HflX or ΔNTE-HflX and evaluated their ability to reassociate with SSU to form 70S (**Figure 3g**). The post-dissociated LSU generated by ΔNTE-HflX is indeed more efficient in forming 70S whereas the post- dissociated LSU generated by full length HflX is not, suggesting a direct role of NTE in disordering H68-71.

This role of NTE in rRNA helical disorder is incongruent with the observation of an ordered H68-71 in 50S-HflX-A, where HflX-NTE is also present. We hypothesize that the pool of ribosomal particles in 50S-HflX-A (5.4 %, fig. S1) represents a small proportion of pre-existing active LSUs rather than a product of HflX-mediated 70S splitting. To test this hypothesis, we solved the cryo-EM structure of a complex of purified *Msm* LSU with full length HflX-GMPPCP for a qualitative comparison. This structure shows distinctly ordered H54a and H68-71 region (fig. S8), which confirms our hypothesis that the 50S-HflX-A complex represents a full-length HflX bound to a pre-dissociated LSU. This also corroborates previous biochemical data for *Eco* HflX ^3^, which suggested a higher affinity for LSUs than 70S ribosomes, and a more pronounced anti-association activity than splitting activity. The structures of *Eco* and *Lmo* 50S-HflX complexes ^3, 5^ were either assembled using LSUs (in *Eco*) or pulled-down with tagged HflX (in *Lmo*), and are likely representative of just the anti-association activity of HflX.

### Structures of the ribosome-HflX complexes in the presence of antibiotics

*Msm* HflX imparts resistance to macrolide-lincosamide antibiotics, with deletion of the *hflx* gene in *Msm* and *Mab* resulting in increased sensitivity to erythromycin, clarithromycin, azithromycin, and clindamycin, but not chloramphenicol ^1^. No structures of HflX-ribosome complexes bound to antibiotics have been reported yet. The *Msm* HD-loop extensively interacts with 23S rRNA bases in and around the PTC that are important in the peptidyl transferase activity (fig. S3b). We superimposed our 50S-HflX-A structure with those of ribosomes bound to these PTC-binding antibiotics ^29–31^, which bind either at the mouth of the nascent polypeptide-exit tunnel (NPET) or at the A-site cleft within the PTC. The superimposition indicates that the *Msm* HflX HD-loop would sterically overlap with these bound antibiotics (fig. S9a).

To investigate antibiotic binding in the presence of HflX on the ribosome, we solved cryo-EM structures for two complexes of *Msm* 70S-HflX-GMPPCP; one with erythromycin (Ery) and other with chloramphenicol (Clm) (fig. S10, S11). Even in the presence of these two different classes of antibiotics, HflX splits the 70S ribosome. Two distinct 50S classes were observed in both cases (fig. S9b-I). One resembled 50S-HflX- A with the HflX-HD-loop extending to the PTC (**Figure 2c, d**; fig. S9b,d,f,h), while the other resembled 50S-HflX-B, with the retracted-bent HflX-HD-loop (**Figure 2c, e**; fig. S9c,e,g,i). Irrespective of whether the HD-loop was extended or bent, all four structures showed distinct densities for antibiotics (fig. S9d,e,h,i). Comparison of the 50S-HflX-B- Ery structure with that of *Eco* 50S-Ery (3J5L)^29^ (see fig. S9j) show similar interactions stabilizing Ery at the NPET of both *Msm* and *Eco* ribosomes (fig. S9j). Specifically, A2281 and A2282 of the 23S rRNA (*Msm* numbering; *Eco* numbers G2057, A2058) interacted with the desosamine hydroxyl at position 5 (P5), while the hydrophobic face of the Ery-lactone ring stacked against the nucleotides A2281 and U2835 (*Msm* numbering; *Eco* numbers G2057, C2611) (fig. S9j).

The conformation of Ery in the 50S-HflX-A-Ery and 50S-HflX-B-Ery structures was nearly identical, suggesting that Ery binds similarly even when the HflX-HD-loop extends to the PTC in in 50S-HflX-A-Ery (**Figure 4a**). Two nucleotides in the vicinity of Ery, however, exhibited significant conformational changes due to presence of the extended HD-loop in 50S-HflX-A-Ery. Specifically, the nucleotide base A2286 rotates 72 degrees and moves 8 Å, while U2730 rotates 33 degrees and moves 2 Å. These conformational changes result in two additional interactions between the Ery sugar hydroxyl at position 3 (P3), U2730, and the Gly197 backbone of HD-loop (**Figure 4a**). In 50S-HflX-A-Ery, the HD- loop avoids a steric overlap with Ery by moving 6.6 Å as compared to its extended conformation in 50S-HflX-A (**Figure 4b**, movie S3). This is accompanied by movements of U2730 (by 17 degrees and 2.0 Å) and A2286 (by 63 degrees and 8.0 Å, **Figure 4b**).

**Figure 4.**
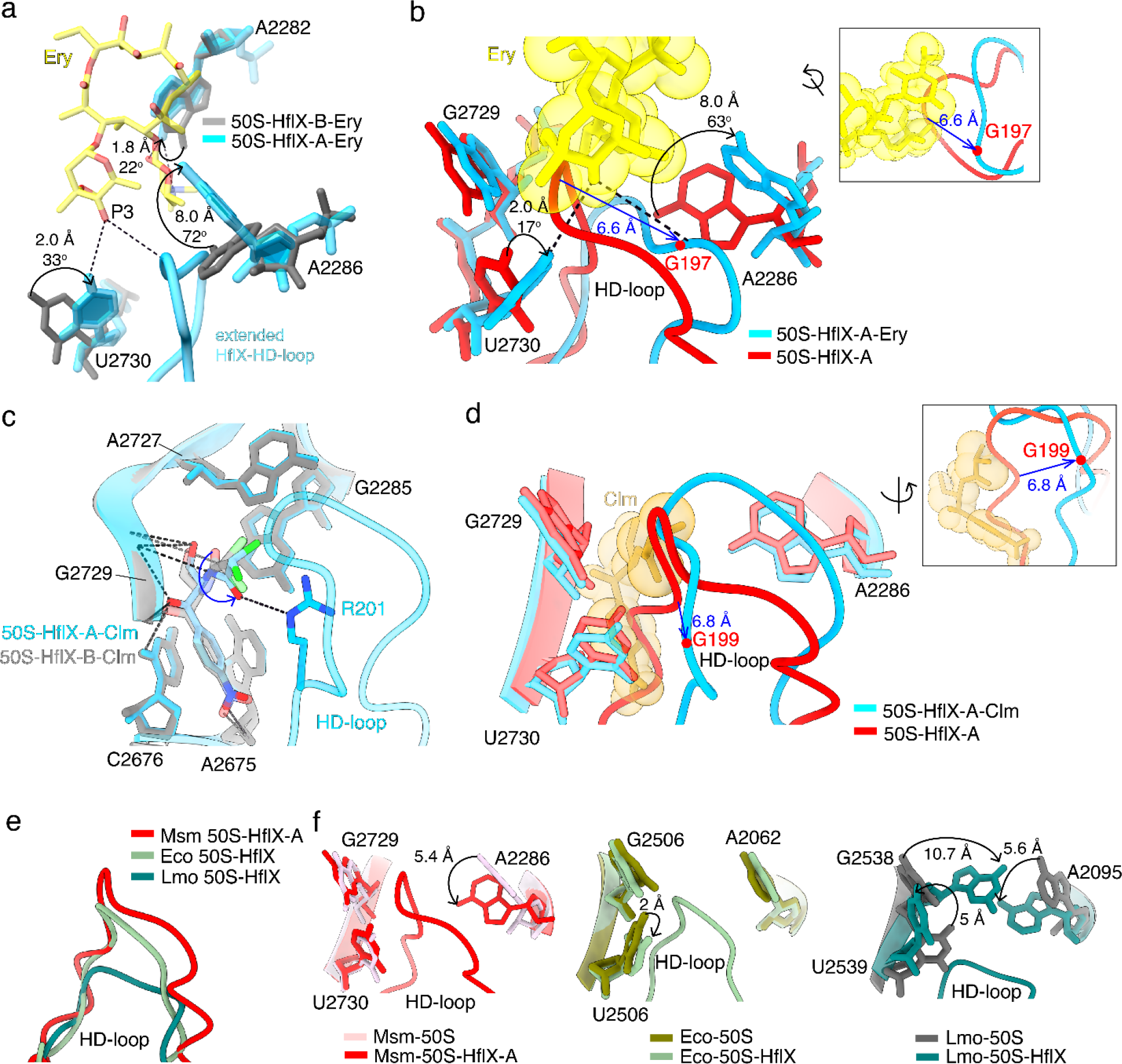
Erythromycin and chloramphenicol binding to the PTC are unaffected by HflX. (a) Superimposition of structures of 50S-HflX erythromycin (Ery) complexes, 50S- HflX-B-Ery (gray) and 50S-HflX-A-Ery (cyan). Interactions of PTC nucleotides and HflX HD-loop to erythromycin (yellow) unique to 50S-HflX-A-Ery are shown. These interactions are due to the rotated state of A2282, A2286 and U2730 in 50S-HflX-A-Ery compared to 50S-HflX-B-Ery. The angular rotations and relative displacement of these nucleotides are indicated (arrows). (**b**) Superimposition of structures of 50S-HflX-A with (cyan) and without (red) erythromycin. The HflX HD-loop is deflected by 6.6 Å in 50S- HflX-A-Ery as compared to that in 50S-HflX-A measured at G197 (marked in red). The inset shows the HD-loop deflection of 6.6 Å in a different view (movie S3). This conformational change in HD-loop is accompanied with rotation (indicated with black arrows) of nucleotide bases U2730 and A2286. (**c**) Superimposition of structures of 50S-HflX chloramphenicol (Clm) complexes, 50S-HflX-B-Clm (gray) and 50S-HflX-A- Clm (cyan). Clm in the 50S-HflX-A-Clm structure additionally interacts with R201 of the HflX HD-loop. (**d**) Superimposition of structures of 50S-HflX-A with (cyan) and without (red) chloramphenicol (gold). The deflection of the HflX HD-loop by 6.8 Å (at residue G199 marked in red) in 50S-HflX-A-Clm compared to 50S-HflX-A is shown. Unlike the erythromycin complexes, this conformational change in HD-loop, does not affect the neighboring ribonucleotides. The inset shows HD-loop deflection of 6.8 Å in a different view (movie S3). (**e**) Superimposition of HflX HD-loops from Msm 50S-HflX-A (red), *Eco* 50S-HflX (light olive green) (PDB:7YLA) and *Lmo* 50S-HflX (teal) (PDB: 8A57) structures, highlighting the difference in length of HD-loops among these species. (**f**) Superimposition of 50S and 50S-HflX-A complex structures from *Msm* (pink, PDB:7Y41 and red), *Eco* (olive green, PDB:7K00 and light olive green, PDB:7YLA) and *Lmo* (gray, PDB:8A63 and teal, PDB:8A57). The conformational changes reported previously in *Lmo* in three PTC nucleotides upon HflX binding is compared with *Eco* and *Msm*.

The extended HD-loop-R201 interacts with Clm in the *Msm* 50S-HflX-A-Clm complex (see **Figure 4c**). The HD-loop in *Msm* 50S-HflX-A-Clm is displaced by 6.8 Å as compared to *Msm* 50S-HflX-A to avoid steric overlap with Clm (see **Figure 4d**, movie S3). The antibiotic binding pocket is very similar in *Msm* 50S-HflX-B-Clm and *Eco*-70S-Clm (PDB: 4V7T) ^32^ but the antibiotic conformation is different (supplementary text, fig. S9k). A comparison between the *Msm* 50S-HflX-B-Clm and Ec-70S-Clm (PDB:4V7T)^32^ structures reveals that while the Clm binding site is the same in the *Msm* structure, there is a significant rotation of the nitro group of the nitrobenzene ring and the amine, along with a displacement of the nitrobenzene ring of the drug (see fig. S9k). Another distinction is noted in the *Msm* structure, where the base of U2730 (U2506 in *Eco*) is not rotated and does not interact with the nitro group of Clm, although interactions with other 23S rRNA nucleotides remain conserved between the two structures (fig. S9k). Further, unlike the Ery-bound structures, the conformations of the 23S rRNA nucleotides U2730 and A2286 remain unperturbed (**Figure 4d**). The Ery and Clm bound structures of 50S-HflX therefore demonstrate that these antibiotics can bind to LSU in the presence of HflX by displacing the extended HD-loop, if necessary.

### Mechanisms of HflX-mediated splitting and antibiotic resistance

Cellular stress caused by increased temperature or antibiotic binding to the NPET can interfere with actively translating 70S ribosomes (**Figure 5a-b**) through peptidyl-tRNA ’drop-off’ to vacate the ribosomes ^33, 34^ or deacylation of the P-tRNA by peptidyl tRNA hydrolases to generate stress-stalled ribosomes ^3, 4^. HflX splits both vacant 70S ribosomes or post- polypeptide released 70S ribosomes in which the P-tRNA is deacylated ^3^ in a GTP- dependent manner ^1, 3^. Our *Msm* 70S-HflX structure (**Figure 5c**, Figure 1a) reveals that the HflX-HD, HflX-GD and HflX-CTD initially anchor to the Sb and adjacent 23S rRNA helices of the 70S. SSU head rotation lets HflX-ND1 bind, likely followed by accommodation of HflX-CTD, which completes HflX accommodation (**Figure 5d)**. This displaces H69, disrupts inter-subunit bridge B2a ^3^, and initiates 70S splitting.

**Figure 5.**
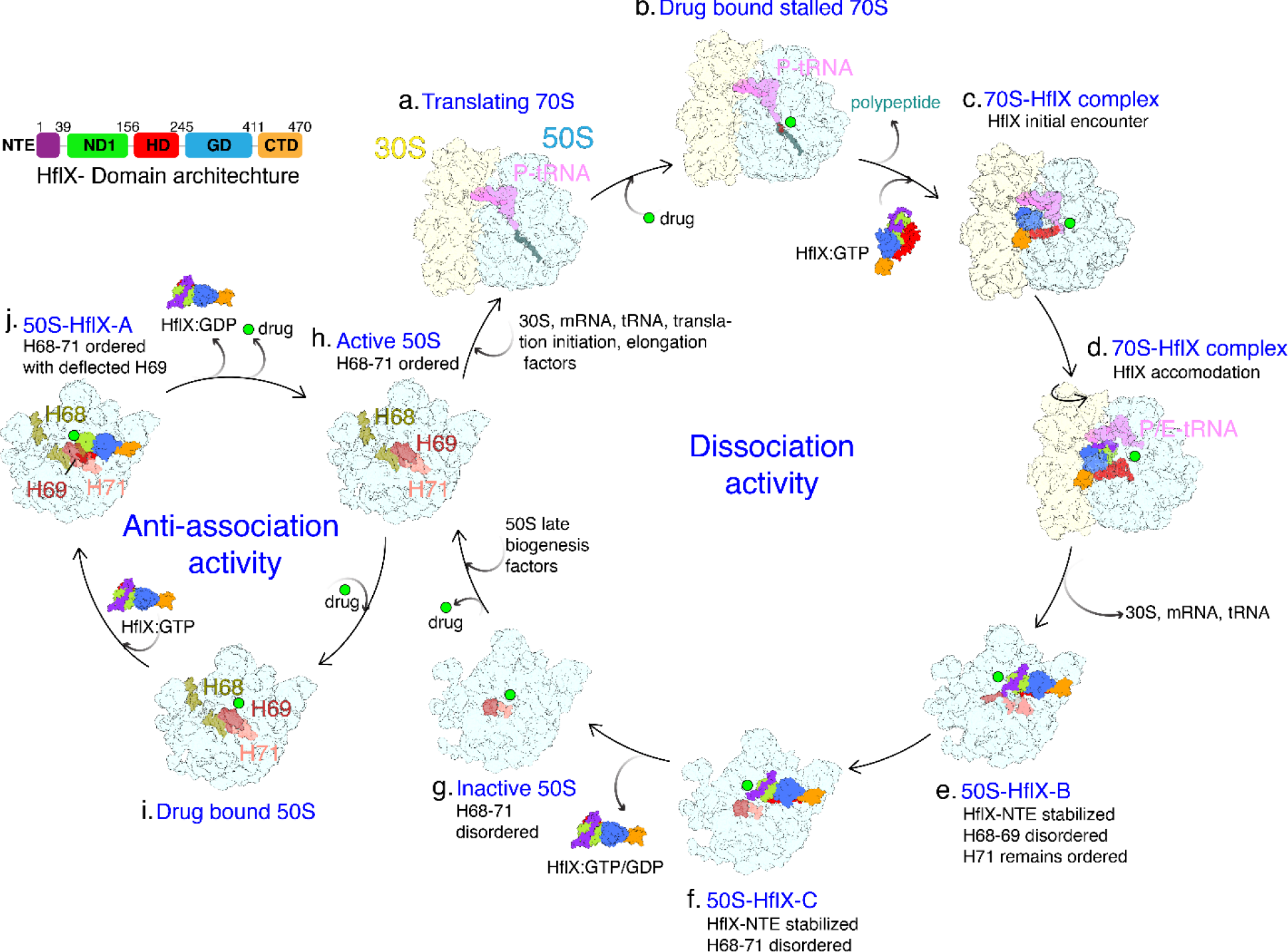
The proposed mechanistic model of the *Msm* HflX-mediated 70S ribosome splitting, antibiotic resistance and anti-association activities. (**a**) An actively translating 70S ribosome with a P-tRNA bound to a polypeptide chain. (**b**) Antibiotic binding would stall translation, leading to release of the polypeptide chain by activity of P-tRNA hydrolases. (**c**) A P tRNA bound 70S ribosome could recruit HflX in its GTP-bound state without the SSU rotation step. At this stage, HflX is in a partially accommodated state via its GD, HD and CTD domains. Interaction of the HD with the PTC ensures that the polypeptide is released. Further, HflX is accommodated along with the drug on the ribosome unlike previous suggestions ^5, 6^ (d) SSU-head rotation ensures transition of the deacylated P-tRNA to the P/E state allowing accommodation of HflX-ND1. (**e**) HflX-NTE stabilization leads to disordering of the 23S rRNA helices H68 and H69 which also facilitates ribosome splitting. (**f**) HflX-NTE further disorders H71 in addition to H68 and H69 before it falls off from the ribosome. (**g**) Inactive 50S LSU with disordered helices H68-71 is retained even after HflX fall-off (fig. S5). (**h**) These inactive LSUs would then require ribosome biogenesis factors to restore their active form (**i**) Antibiotics targeting the PTC can bind to pre-dissociated LSUs. (**j**) HflX can bind to pre-dissociated LSUs without displacing the bound drug molecule. HflX binding to LSU also displaces H69, making it incompatible for association with SSU, which is how HflX also acts as an anti-association factor. (**h**) GTP hydrolysis leads to HflX fall-off from a pre-dissociated LSU. Displacement of the bound drug molecule from otherwise active LSU would allow its entry into elongation phase (**a**).

Stabilization of HflX-NTE through interactions with uL16 and H38 accompanies H68-69 disorder (**Figure 5e**), with a smaller extent of additional H71 disorder (**Figure 5f**). HflX- NTE is required for this H68-71 disorder, as indicated by the structure of the 50S-ΔNTE- HflX complex, which does not show such disorder. The LSUs split from 70S ribosomes by *Msm* HflX are arrested in an inactive conformation through 23S rRNA disorder, even after it unbinds (**Figure 5g**; Figure 3g, fig. S5). This is shown by the LSU structure not bound to HflX still showing a disordered H68-71 region (**Figure 5f**). These disordered 23S RNA helices are involved in multiple inter-subunit bridges ^12, 14, 23–25^ (Figure 3d-f), and their disorder is incompatible with SSU and LSU either remaining associated or re- associating. We biochemically established the role of NTE in disordering H68-71 by showing that full-length HflX-split LSUs have significantly reduced ability to reassociate with 30S ribosomal subunits to form 70S ribosome as compared to *Mab* ΔNTE-HflX-split LSUs (Figure 3g).

Our inactive LSU structures resemble late intermediates of LSU biogenesis, where H68- 71 are still not structured ^26–28^. Biogenesis factors are needed to fold these 23S RNA helices in their active conformation during ribosome assembly (**Figure 5h**). Accumulation of inactive LSUs, with H68 in an altered conformation, has been observed in *Msm* during the stationary phase ^35^. In response to hypoxic stress in *Mbo*, HflX modulates bacterial growth rate and facilitates entry into a non-replicating dormant state ^4^. The *Mbo* HflX, which also has a longer NTE, could similarly generate a pool of inactive LSUs by 70S ribosome splitting and 23S RNA helical disorder that leads to dormancy.

Another structurally characterized HflX homolog is the human mitochondrial GTPB6 ^36^. The cryo-EM structure of its assembly intermediate with the human mitochondrial 39S large ribosomal subunit resembles the 50S-HflX-A state, with its HD reaching the PTC in an extended state, while its NTE is disordered. Recent time-resolved cryo-EM (TRCEM) studies on *Eco* and *Lmo* capture the initial, intermediate, and final states during 70S splitting by HflX, but do not report disordering of the rRNA helices in the resulting LSU ^10, 11^ (supplementary text, fig. S12). Both *Eco* and *Lmo* HflX lack an NTE and their lack of persistent 23S RNA disorder is consistent with our finding that NTE is necessary to disorder large subunit ribosomal RNA helices during splitting. While the human mitochondrial GTPB6 does carry a long NTE, the H68-71 in the mitoribosome is significantly shorter (only 54 nucleotides *versus* 146 nucleotides in *Msm*). This raises two possibilities: either the NTE disorders these helices but they are capable of readily refolding due to their shorter size, or both the NTE and H54a play a role in the process, where the NTE mediates the disorder of H68-71 while H54a helps in sustenance of the disordered state.

In *Eco*, HflX preferentially binds to LSUs, inhibiting their association with empty 30S subunits, but an HflX-bound LSU can still bind a 30S pre-initiation complex ^3^ to form 70S ribosome. GTP hydrolysis is also required for HflX release from *Eco* LSUs ^3^. When we prepare a complex of the *Msm* 70S-HflX, HflX shows a preferential binding to entire population (∼5 %) of the pre-dissociated LSUs present as a contaminant in the 70S sample (Figure 1c, d) to form the 50S-HflX-A complex with H68-71 ordered and the HflX- NTE disordered. In this complex, HflX forms extensive interactions with the neighboring 23S rRNA. Release of HflX from the 50S-HflX-A complex would require GTP hydrolysis, which could be stimulated by a 30S initiation complex. Fewer interactions destabilize the altered conformation of HflX on LSUs formed by 70S splitting (50S-HflX-B and 50S-HflX- C), which explains HflX displaying a higher affinity for pre-dissociated LSUs than for the LSUs derived from 70S splitting. It could also account for the spontaneous release of GMPPCP-bound HflX from LSUs obtained from 70S splitting. Thus, when HflX is overexpressed under stress conditions, it could bind to pre-dissociated LSUs, acting as an anti-association factor that prevents premature formation of 70S ribosomes.

*Msm* and *Mab* HflX impart significant resistance to macrolide-lincosamide antibiotics ^1^, which bind to actively translating ribosomes and result in the accumulation of stalled 70S ribosomes (**Figure 5b)** or free LSUs (**Figure 5i**). In *Eco*, a Type I resistance mechanism for Clm in the presence of HflX was suggested, where HflX displaces the antibiotic or prevents its binding ^6^. In *Lmo*, a Type II resistance mechanism was proposed, where HflX binding allosterically modifies the antibiotic binding pocket to prevent or reduce antibiotic binding ^5^. Our 50S-HflX-A structure shows that the longer *Msm* HD-loop does extends deeper into the PTC compared to *Eco* (PDB ID: 7YLA) and *Lmo* HflX (PDB ID: 8A57) (Figure 4e), which agrees with a Type I mechanism at first glance. However, our structures of 50S-HflX with two antibiotics, Ery or Clm, show that both Ery and Clm can bind to the *Msm* ribosome irrespective of whether the conformation of the HflX-HD-loop is extended or retractive-bent (Figure 4a-d). Even when the HD-loop extends to the PTC, it adjusts around both Ery and Clm (Figure 4b, d, movie S3) instead of displacing them. Given the known flexibility of the HflX HD-loop across orthologs, we expect similar behavior in other organisms ^5, 10^.

Certain PTC nucleotides, U2730 and A2286 (*Msm* numbering; U2506 and A2062 in *Eco*; U2539 and A2095 in *Lmo*), do undergo concomitant conformational changes with HD- loop motion (Figure 4f). Our antibiotic-bound structures show that these conformational changes of the binding pocket do not prevent antibiotic binding, and may even contribute to additional antibiotic stabilization (Figure 4c, d). Neither a Type I nor a Type II mechanism is therefore supported by our structural data for *Msm* HflX-mediated antibiotic resistance. This is also consistent with the flexible nature of the HflX HD-loop across orthologs, which suggests that it may similarly adjust to antibiotic presence in other organisms as well.

An alternative mechanism of mycobacterial HflX antibiotic resistance, which agrees with our structural data, is that HflX dissociates antibiotic-stalled 70S ribosomes to a pool of inactive LSUs that sequester bound antibiotics. Subsequently, ribosome biogenesis or other downstream factors may recycle these LSUs for their return to active protein synthesis (**Figure 5 g-h**), with a slower release of antibiotic that is less detrimental to the cell. A possible reason why *Msm* HflX-mediated antibiotic resistance is not seen for certain antibiotics, such as Clm, is that they may rely on non-HflX-mediated rescue of stalled ribosomes. In contrast, the characteristic induction of peptidyl tRNA dropoff by the macrolide-lincosamide class of antibiotics ^33, 34^ serves as an ideal substrate for HflX-mediated rescue of stalled 70S ribosome.

## Methods and Materials

### Construction of recombinant plasmids, overexpression and purification of recombinant HflX and purification of *Msm* ribosomes

*Msm* HflX and ΔNTE HflX were purified from *Eco BL21 (DE3)pLys* as described previously for *Msm* HflX ^1^. Ribosomes were purified from *Msm* wild type mc^2^155 as described previously ^1^.

### *In vitro* reconstitution of the 70S-HflX complex

Purified *Msm* 70S ribosomes (300 nM) were mixed with either full-length HflX or ΔNTE HflX proteins at final concentrations of 3 µM (for a 10-fold molar excess) or 6 µM (for a 20-fold molar excess), together with 200 µM puromycin and 1 mM GMPPCP in HMA-10 buffer (containing 20 mM HEPES-K at pH 7.5, 30 mM ammonium chloride, 10 mM magnesium chloride, and 5 mM ß- mercaptoethanol). Each reaction mixture was incubated at 37°C for 30 minutes. To prepare the *Msm* 70S-HflX-antibiotics complexes, 300 nM of 70S ribosomes were first incubated on ice for 30 minutes with 100 μM of the antibiotics. Subsequently, they were combined with 6 μM of full-length HflX, 200 µM puromycin and 1 mM GMPPCP and further incubated at 37°C for an additional 30 minutes.

### Re-association of post-dissociated LSU

The *Mab* cell lysates were solubilized in HMA6/30 [20 mM HEPES (pH 7.5), 6 mM MgCl2, 30 mM NH4Cl, 5 mM 2-Mercaptoethanol] and clarified by centrifugation at 30,000 × *g* for 30 min, 4 °C. Clarified lysates were treated with Turbo DNase (5U/mL) and puromycin (2000-fold molar excess) for 3h at 37 °C. A crude ribosome pellet was obtained by centrifugation for 2h at 200,000 × g at 4 °C (70Ti rotor, Beckman Coulter) and solubilized in HMA6/30 for 16h at 4°C. To split the 70S ribosomes, 5-fold molar excess HflX or ΔNTE-HflX was added in presence of GTP (1mM) and incubated at 37 °C for 2.5h. The ribosomal pellets were obtained by passing the reaction mixture through a 15% sucrose cushion by centrifugation at 200,000 × g (70Ti rotor) for 16h at 4 °C, and were solubilized in HMA6/30, overlaid onto a 10 - 40% sucrose gradient (HMA6/30), centrifuged at 100,000 × g at 4 °C for 18h (SW28 rotor, Beckman Coulter), and fractionated using Brandel gradient fractionator. The LSU fractions were collected, pelleted, and solubilized in HMA6/30.

The SSU was prepared by solubilizing a crude preparation of ribosomes as described above, but in HMA1/30 [20 mM HEPES (pH 7.5), 1 mM MgCl2, 30 mM NH4Cl, 5 mM 2- Mercaptoethanol]. The ribosomal subunits were separated through a 10 - 40% sucrose gradient (HMA1/30) and SSU fractions were collected, pelleted, and solubilized in HMA6/30. The re-association of post-split LSU with SSU was studied by incubating 0.2 μM of each subunit in a 50 μl final volume of HMA8/30 [20 mM HEPES (pH 7.5), 8 mM MgCl2, 30 mM NH4Cl, 5 mM 2-Mercaptoethanol] at 37 °C for 1h, and sucrose gradient profile was obtained on Brandel gradient fractionator.

### Cryo-electron microscopy and image processing

Quantifoil R1.2/1.3 holey carbon copper grids, pre-coated with a 2 nm carbon layer, were further coated with a continuous carbon layer (∼50 nm thick). These grids underwent a 30-second glow-discharge treatment in a plasma sterilizer. For sample vitrification Vitrobot IV (FEI) was used. Four microliters (4 µL) of the respective ribosome-HflX complex were applied to the grids and incubated for 15 seconds at 4°C under 100% humidity, followed by blotting for 4 seconds to remove excess sample, before plunge freezing the grids in liquid ethane. Cryo-EM micrographs were acquired using a Titan Krios electron microscope operating at 300 keV, equipped with a K3 direct electron detecting camera (Gatan). The collected datasets were processed in cryoSPARC v3 ^37^.

The data collection parameters, detailed processing strategy for the individual datasets and model refinement parameters are provided below: *70S-full length HflX complex prepared with 10-fold molar excess of HflX:* The micrographs were captured at a nominal magnification of 81,000x, with a pixel size of 1.08 Å and a total dose of 51.22e-/Å2. The defocus range varied between -1.8 and -2.5 µm. We performed alignment on 4,766 movies using patch-motion correction and estimated their CTF parameters. Subsequently, micrographs were manually curated based on the CTF results, resulting in the removal of poor-quality micrographs and leaving us with 4,668 micrographs. From these micrographs, a total of 1,529,798 particles were picked using a template-based particle picking approach. Template-based particle picking was performed using a reference template generated from a low-pass filtered map of the Msm 70S ribosome ^13^. These particles underwent several rounds of 2D classification to eliminate non-ribosomal particles (fig. S1). After 2D classification, 1,051,589 particles were retained and subjected to further processing through heterogeneous refinement, resulting in four distinct 3D classes. Among these classes, three represented 70S ribosomes (994,155 particles), while one class, consisting of 57,434 particles, corresponded to the 50S-HflX complex, State A. This 50S-HflX-A class was refined to an overall resolution of 3.14 Å. Additionally, one of the 70S classes with 537,249 particles, displayed a weak density for HflX and P and E-site tRNAs. This class was further classified, leading to the isolation of a 70S class with E-site tRNA and HflX (48,443 particles). This class was refined to an overall resolution of 3.26 Å.

*70S-full length HflX complex prepared with 20-fold molar excess of HflX:*Micrographs were collected at a magnification of 64,000x, with a pixel size 1.07 Å, total dose of 51.85 e-/Å2 and a defocus range of -1.0 to -2.5 µm. 8,934 movies were aligned using patch-motion correction and their CTF parameters estimated. 8,816 micrographs remaining after manual curation were used for template-based particle picking (as discussed above). 3,123,836 particles obtained from particle picking was manually curated and 3,053,229 particles were used for 2D-classification (fig. S5). After several rounds of 2D- classifications to eliminate non-ribosomal particles 1,685,480 particles were retained. These particles were classified using 3D heterogenous refinement. Except for a small class of the 70S ribosome with 66,901 particles, majority of the dataset with 1,357,294 particles were LSUs with a weak density for HflX. All 50S LSU classes were recombined and classified further to obtain a homogenous set of 50S subunits with 1,223,631 particles. 3D-variability analysis was carried out with these particles using a mask encompassing the HflX binding region on LSU. Based on the 3D-variability analysis the particles were distributed into ten 3D classes. A significant population of LSU (787,017 particles) were found to be empty (without HflX). From the HflX-bound LSU classes, we refined two unique classes, one with disordered H68-69 but an ordered H71 (State B) and the other with disordered H68-71 (State C). We also refined a LSU class with no HflX but disordered H68-71. The overall resolutions of these three maps were 3.09 Å (State B), 2.96 Å (State C) and 3.0 Å (LSU without HflX).

*70S-ΔNTE HflX complex prepared with 20-fold molar excess of ΔNTE HflX:* The micrographs were captured at a magnification of 81,000x, with a pixel size of 0.846 Å, a total dose of 60.64 e-/Å2, and a defocus range spanning from -0.8 to -2.5 µm. A total of 8,700 micrographs were subjected to patch-motion correction for alignment, and their CTF parameters were subsequently estimated. After a manual curation process based on the CTF parameters, 8,362 high-quality micrographs were retained. These selected micrographs were utilized for template-based particle picking that resulted in 2,353,863 particles. Following a thorough manual inspection of the particles, 417,953 particles were retained for further processing and subjected to 2D-classification to eliminate non-ribosomal classes. This process yielded 273,134 ribosomal particles (fig. S7). Heterogeneous 3D refinement was then employed to classify these particles, leading to the separation of a LSU class consisting of 91,763 particles and a 70S class comprising 181,371 particles. It is noteworthy that the LSU class exhibited a weak density for HflX. Subsequently, a 3D-variability analysis was conducted on these particles using a mask encompassing the HflX binding site on LSU. As a result of this analysis, a class with a relatively homogenous population with 27,550 particles of the 50S-HflX complex was obtained and refined to an overall resolution of 3.4 Å.

*70S-full length HflX complex prepared with 20-fold molar excess of HflX and Erythromycin:* Micrographs were captured at a magnification of 81,000x, with a pixel size of 0.846 Å, a total dose of 62.87 e-/Å2, and a defocus range of -1.0 to -1.6 µm. Initially, 12,145 movies were aligned using patch-motion correction, and their CTF parameters were estimated. Following manual curation, 11,594 micrographs were selected for template-based particle picking. Subsequently, 3,113,258 particles obtained from particle picking underwent manual curation, resulting in 2,175,541 particles for 2D-classification (fig. S10). After several rounds of 2D-classifications aimed at eliminating non-ribosomal particles, 532,420 particles were retained. These particles underwent classification using Ab-initio reconstructions (fig. S10) to exclude non-ribosomal particles and those with preferred orientations. The Ab-initio reconstructions revealed only 50S classes, suggesting that the majority of the 70S ribosomal subunits had split into 50S and 30S components. Subsequently, 366,321 LSU particles were recovered from the Ab-initio reconstructions and subjected to masked 3D classification. The mask encompassed H68- 71, H54a, and HflX. This classification yielded three distinct classes: 50S-HflX-A-Ery (46,709 particles), 50S-HflX-B-Ery (113,394 particles), and empty LSU (206,218 particles). The first two classes were refined to resolutions of 2.8 Å and 2.63 Å, respectively (fig. S10).

*70S-full length HflX complex prepared with 20-fold molar excess of HflX and Chloramphenicol:* Micrographs were captured at a magnification of 105,000x, with a pixel size of 0.8443 Å, a total dose of 70.14 e-/Å2, and a defocus range of -0.8 to -2.5 µm. Initially, 14,374 movies underwent patch-motion correction and had their CTF parameters estimated. Following manual curation, 13,644 micrographs were chosen for template-based particle picking. Subsequently, 3,809,121 particles obtained from particle picking underwent manual curation, resulting in 2,563,191 particles for 2D-classification (fig. S11). After several rounds of 2D-classification aimed at eliminating non-ribosomal particles, 542,078 particles were retained. These particles were classified using Ab-initio reconstructions (fig. S11) to exclude non-ribosomal particles and those with preferred orientations. The Ab-initio reconstructions revealed only 50S classes, suggesting that the majority of the 70S ribosomal subunits had split into 50S and 30S components. Subsequently, 427,915 LSU particles were recovered from the Ab-initio reconstructions and subjected to masked 3D classification. The mask included H68-71, H54a, and HflX. This classification resulted in three distinct classes: 50S-HflX-A-Clm (65,766 particles), 50S-HflX-B-Clm (116,149 particles), and empty LSU (246,000 particles). The first two classes were refined to resolutions of 3.3 Å and 3.25 Å, respectively (fig. S11).

### Model building

Coordinates of our previously published *Msm* C- ribosome structure (PDB:6DZI, 70S or LSU) was docked as rigid bodies into the respective maps using Chimera 1.14 ^38^. The models were modified based on the cryo-EM densities, for optimal fitting, in COOT ^39^. The models were subsequently refined in PHENIX 1.14 ^40^. The *Msm* HflX predicted structure was available in the Alphafold Protein Structure database ^41, 42^. This structure was docked into the corresponding maps, locally readjusted with COOT ^39^ on the cryo-EM density and finally refined using PHENIX 1.14 ^40^. The NTE of HflX was built manually since its conformation did not entirely match the predicted Alphafold model. The RNA and protein geometries of the models are validated using MolProbity. The overall statistics of EM reconstruction and refinement of the molecular models are listed in tables S1-S3. The structure figures in the manuscript are generated using Chimera X 1.0 ^43–45^ and Chimera 1.14 ^38^.

## Supporting information

an animation showing conformational changes in mycobacterial HflX on the 50S ribosome

an animation showing stepwise disordering 23S rRNA helices upon mycobacterial HflX binding to the 70S ribosome

an animation showing how the insertion loop in mycobacterial adjusts its structure around the 50S-bound antibiotics

## Acknowledgements

This work was supported by grants to PG and RKA (NIH: R01 Al155473) and, in part, by a grant to RKA (NIH: R01GM61576). RKA also acknowledges support to his lab through NIH R01 grants AI132422 and GM139277. Authors acknowledge Wadsworth Center’s and New York Structural Biology Center’s (NYSBC’s) 3D-EM facilities. Wadsworth Center is a contributing member of NYSBC, whose EM facility is supported by the Simons Foundation (SF349247).

## Author Contributions

PG and RKA conceived this study. AK, KH-H purified Msm ribosomes, created Msm and Mab full length HflX and ΔNTE-HflX constructs and performed biochemical characterization. SM, RKK, and RKA designed structural experiments. RKK prepared ribosome-HflX complexes, and RKK, MRS, and NKB performed initial structural work with full length HflX. SRM and NKB collected cryo-EM data and performed initial reconstructions of multiple samples from data collected on the in-house cryo-electron microscope. SM purified ΔNTE-HflX, prepared multiple ribosome-HflX complexes and all the ribosome-HflX-antibiotics complexes, performed detailed image processing, performed molecular modeling and functional analyses of all complexes. NKB and RKA contributed to structural analyses. NKB generated movies with inputs from SM and RKA. SM and RKA wrote the manuscript, with help from MRS, NKB and PG. All authors read and approved the manuscript.

## Competing Interests

The authors declare no competing interests.

## Data Deposition

Accession codes for the cryo-EM density maps and their corresponding atomic coordinates for the following nine complexes that have been deposited in the Electron Microscopy Data Bank [https://www.ebi.ac.uk/pdbe/entry/emdb/] and the Protein Data Bank [https://www.rcsb.org/structure/], respectively, are 70S-HflX complex (EMD-43267 and PDB ID-8VIO), 50S-HflX-A complex (EMD-43294 and PDB ID-8VK0), 50S-HflX-B complex (EMD-43305 and PDB ID-8VK7), 50S-HflX-C complex (EMD-43317 and PDB ID-8VKI), 50S-ΔNTE-HflX complex (EMD-43333 and PDB ID-8VKW), 50S-HflX-A-Ery complex (EMD-43476 and PDB ID-8VR4), 50S-HflX-B-Ery complex (EMD-43409 and PDB ID-8VPK), 50S-HflX-A-Clm complex (EMD-43484 and PDB ID-8VRL), and 50S- HflX-B-Clm complex (EMD-43477 and PDB ID-8VR8). Accession codes for the cryo-EM maps of three complexes that were obtained as control for qualitative comparison are also deposited in the Electron Microscopy Data Bank [https://www.ebi.ac.uk/pdbe/entry/emdb/] are 70S-P-tRNA complex (EMD-43791), Inactive-50S (EMD-43778), and pre-dissociated 50S-HflX complex (EMD-44044).

List of Supplementary Materials

Figs. S1 to S13 Tables S1 to S3 References (1-3) Movies S1-S3

## Notes

### Competing Interest Statement

The authors have declared no competing interest.

