## Supplementary text, figures, and tables for "Drug resistance through ribosome splitting and rRNA disordering in mycobacteria"

***Comparison with results of recent time-resolved cryo-EM studies in Eco and Lmo HflX.*** After completion of this study, two time-resolved cryo-EM (TRCEM) studies on the mechanism behind HflX-mediated splitting 70S ribosomes was published[1,2]. Three intermediate states of the 70S-HflX complex at TRCEM time points 10, 25, and 140 milliseconds were captured. It was found that accommodation of HflX accompanied with rotation of the 30S ribosomal subunit (SSU) with respect to the 50S ribosomal subunit (LSU), disrupts a series of inter-subunit bridges that results in the splitting of the 70S ribosome.

By superimposing our *Msm* 70S-HflX structure with the three *Eco* 70S-HflX intermediates (fig. S12 a-f), we find that the SSU-body in our structure resembles the second intermediate (25 ms, *Eco* i70S-HflX-II, fig. S12b), while the head lies at an intermediate position between the second and third intermediate states (10 ms, *Eco* i70S-HflX-I and 25 ms, *Eco* i70S-HflX-II, fig. S12a,b). The positions of ribosomal proteins uL2 and bS6 in our structure closely resemble intermediate II (fig. S12e). Thus, our structure shows an overall resemblance to the *Eco* i70S-HflX-II structure (fig. S12b, e).

The conformation of HflX and the LSU-stalk base (Sb) differs in our structure compared to the intermediate II (fig. S12g). Specifically, in our structure HflX-CTD primarily interacts with the 23S rRNA helix (H95) (**Figure 1g**; fig. S12g and 12g-2), whereas H43 of the Sb interacts with the GD (**Figure 1g**; fig. S12g). In contrast, in the *Eco* intermediate II structure, it was suggested that *Eco* HflX-CTD interacts with the Sb. The cryo-EM density (as well as the corresponding molecular model) for uL11 are not provided (fig. S12g-1) and those of H43-44 and HflX-CTD are incipient (fig. S12g-1).

A comparison of the three *Eco* 70S intermediates reveals that the movement of HflX-CTD does not follow a smooth pattern (fig. S12h). First, it moves by 15.4 Å towards the Sb from P1-P2, deflecting the Sb by 8.5 Å (p1-p2) (fig. S12h), followed by its deflection away from the Sb by 19.6 Å, as the Sb returns to its initial position (p3=p1) (fig. S12h). Finally, it moves by 12.8 Å to P4 in the *Eco* 50S-HflX complex without any movement of the Sb (p4=p1) (fig. S12h). The cryo-EM densities for the *Eco* HflX and Sb of the ribosome are relatively weak in these intermediates. In our *Msm* 70S-HflX structure, we observe a smooth movement of the Sb and HflX-CTD. The Sb interacts with HflX-GD (likely to check the nucleotide-bound state of the factor) in the 70S complex. The HflX-CTD then moves by 18.6 Å towards the Sb, which itself moves by 13.3 Å for this interaction to occur (fig. S12i) in the 50S-HflX complex.

Another TRCEM study involving *Lmo* HflX and *Listeria innocua* (*Lin*)70S ribosomes identifies three distinct HflX-bound 70S intermediates, with one of these intermediates also engaging a hibernation promoting factor (HPF)[2]. Through a comparative analysis of small changes in the buried surface area at inter-subunit bridges between HflX-unbound and bound ribosomal intermediates, the authors propose that the displacement of H69 alone does not disrupt bridge B2a. Instead, the displacement of H69, in conjunction with 30S rotation, induces movements in the 30S platform domain, leading to the disruption of multiple inter-subunit bridges and consequent 70S splitting. A comparison with variations in buried surface area of the inter-subunit bridges inherent to ribosomal dynamics during protein synthesis was not made.

Both TRCEM studies reported with *Eco* and *Lmo* HflX do not report the 23S rRNA helical disorder associated with HflX-mediated splitting of 70S ribosome found in our study. There are two possible explanations: (i) disordering of the rRNA helices is specific to mycobacterial HflX or other HflX orthologs with an NTE; or (ii) this rRNA helical disordering is a transient part of all HflX-mediated 70S splitting, and the presence of the NTE in mycobacterial HflX prevents restoration of the ordered states of these helices. In the latter case, intermediates with disordered rRNA helices in *Eco, Lmo* and *Lin* could be occurring in intermediate time points not sampled in the TRCEM studies.

### Supplementary Figures

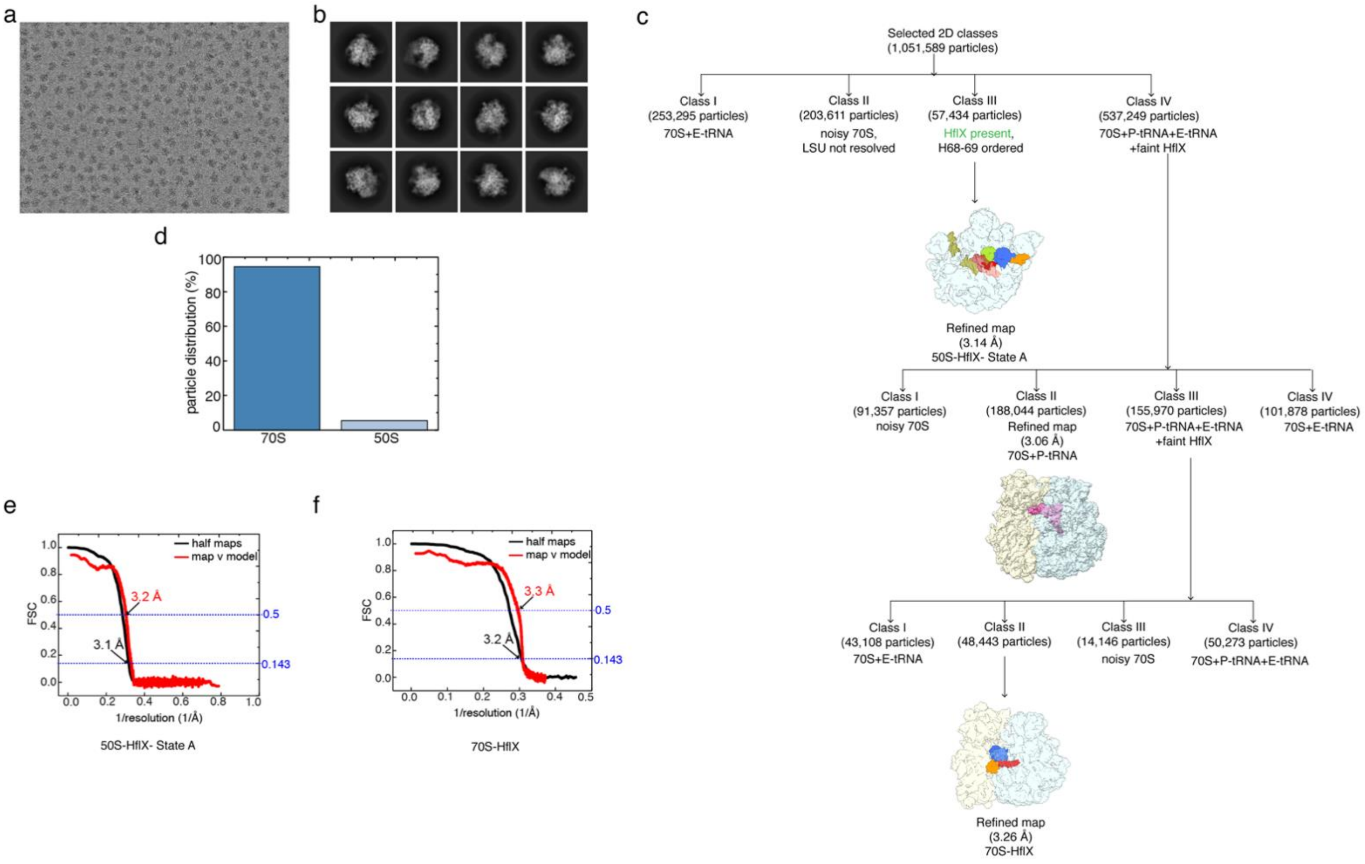


**Supplementary Figure 1. Image processing of *M. smegmatis* 70S-HflX complex prepared with lower concentration of HflX:GMPPNP**. (**a**) a representative micrograph from the cryo-EM dataset. (**b**) Representative 2D-class averages selected for subsequent data-processing. (**c**) Flowchart showing the details of 3D classifications and refinements performed on the selected particles. The final two maps used for model building and interpretations (in main Figure 1) are displayed. (**d**) Plot comparing the percentages of 70S and 50S ribosomal subunit obtained. Gold-standard FSC (black) overlaid with the map-to-model FSC (red) for (**e**) the 50S-HflX-State A and (**f**) the 70S-HflX maps.

**Supplementary Figure 2.**

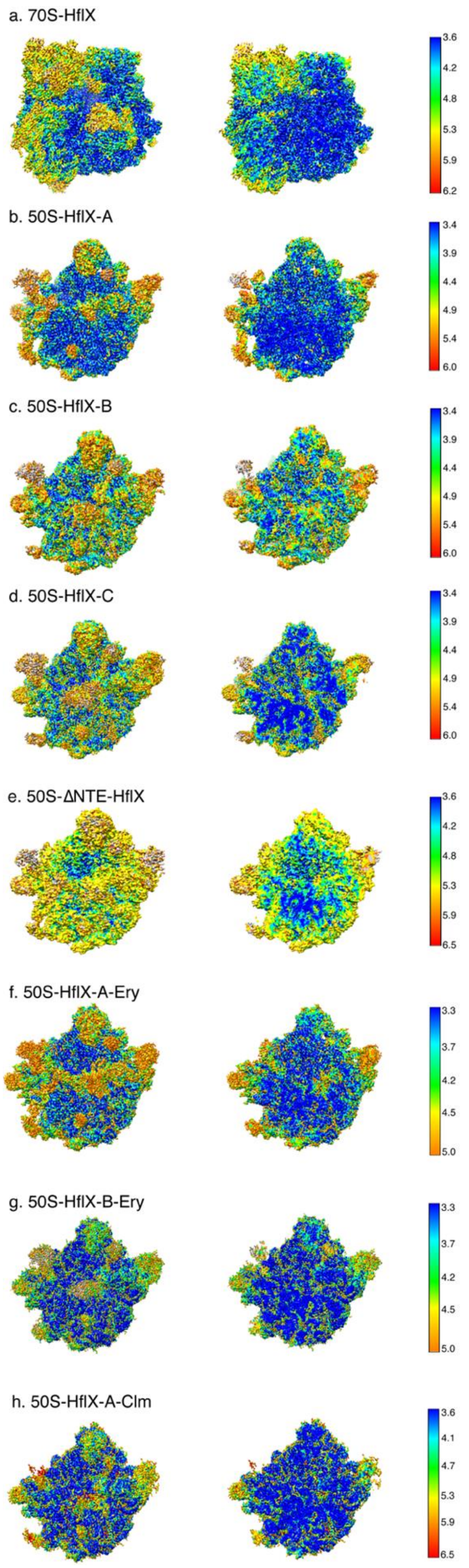

**Supplementary Figure 2. Local resolution of the cryo-EM maps obtained for the ribosome-HflX complexes.** Local resolution of the maps was estimated using Resmap[3]. The densities are colored based on the local resolution and the color keys are provided at the end of each row. The first column of (**a-h**) represents the complete map while the second column shows slices through the same maps.

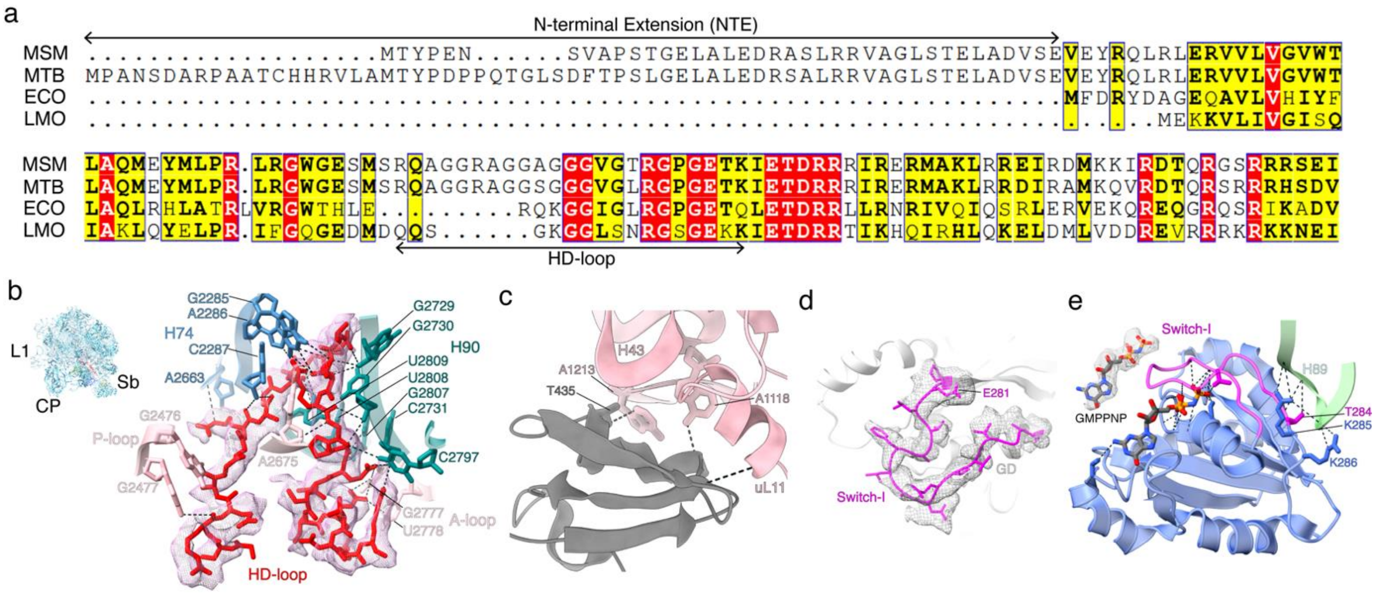


**Supplementary Figure 3.** (**a**) Sequence alignment of the N-terminal extension (NTE) and the HD-loop region of representative HflX homologs from *Mycobacterium smegmatis* (MSM), *Mycobacterium tuberculosis* (MTB), *Escherichia coli* (ECO) and *Listeria monocytogenes* (LMO). (**b**) The density corresponding to the loop connecting the two α-helices (HD-loop) in the 50S-HflX-A structure, along with its interactions with the PTC components, P-loop, A-loop, H74 and H90 are shown. (**c**) Interactions of the CTD in 50S-HflX-A with components of the L7/L12 stalk base, uL11 and H43. (**d**) Density corresponding to the Switch-I (magenta) of 70S-HflX-GD. (**e**) Molecular interactions of the 50S-HflX-A GD with H89. The Switch I (magenta) is ordered and is in the “ON” state. Switch I T284, K285 and K286 interact with H89. The cryo-EM density of the ligand, GMPPNP is also displayed.

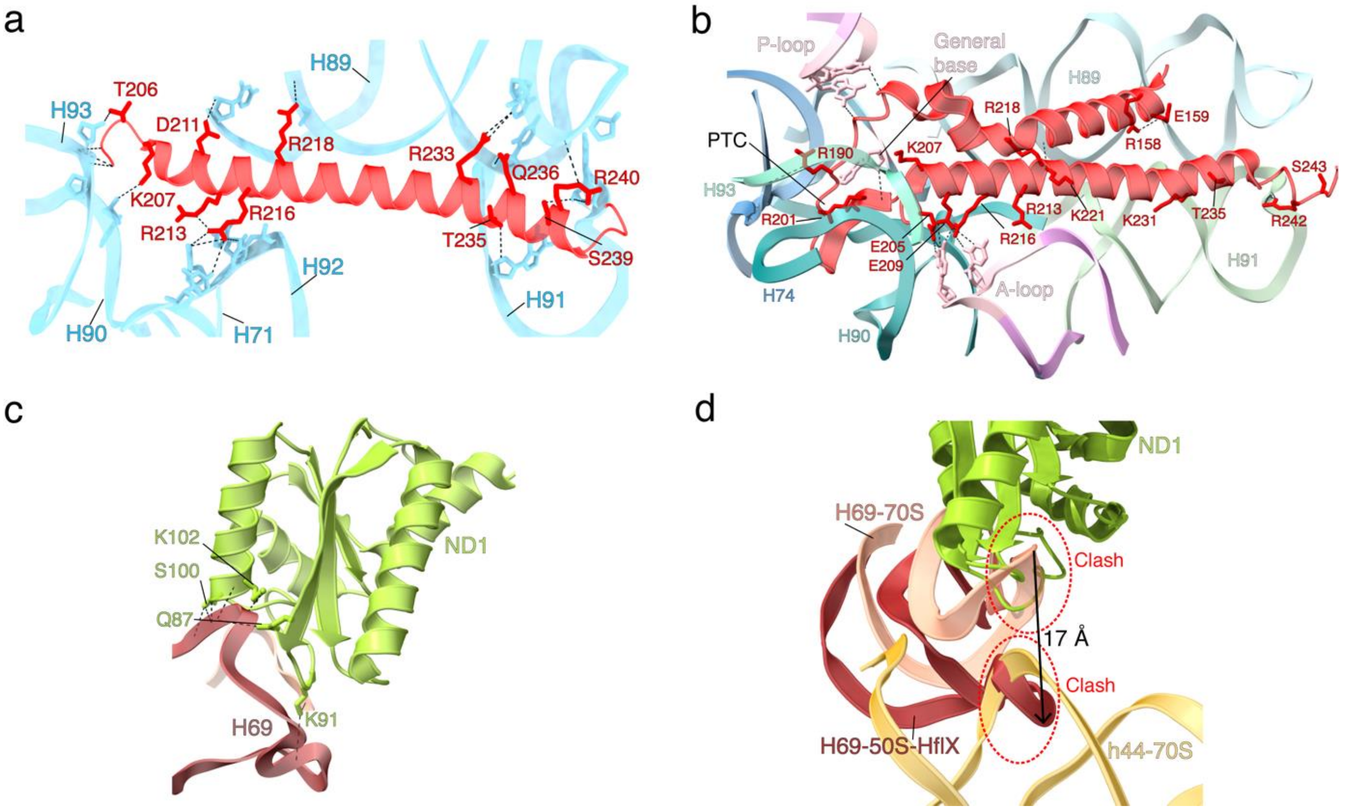


**Supplementary Figure 4.** (**a**) Molecular interactions of the HflX-HD with the 23S rRNA helices, H71 and H89-93 in the *Msm* 70S-HflX complex. (**b**) Molecular interactions of the HD (red) with the neighboring 23S rRNA helices and loops, H74 (steel blue), H89 (light green), H90 (teal), H91 (dark sea green), H93 (medium aquamarine), and components of the peptidyl transferase center (PTC), P-loop (pink) and A-loop (pink) in the 50S-HflX-A complex. Interaction with the general base for the peptidyl transfer reaction, A2675, is also shown. The loop connecting the two α-helices extend into the PTC. (**c**) Molecular interactions of ND1 with H69 in the 50S-HflX-A complex. (**d**) Superimposition of the 50S-HflX-A structure with 70S monosome structure (PDB:6DZI). H69 in the 50S-HflX-A (brown) is displaced by 17 Å, as compared to its position in the 70S monosome (light salmon). This displaced conformation of H69 in the 50S-HflX-A would directly clash with h44 in the 70S monosome, thereby preventing association of the HflX-bound 50S subunit to the 30S subunit.

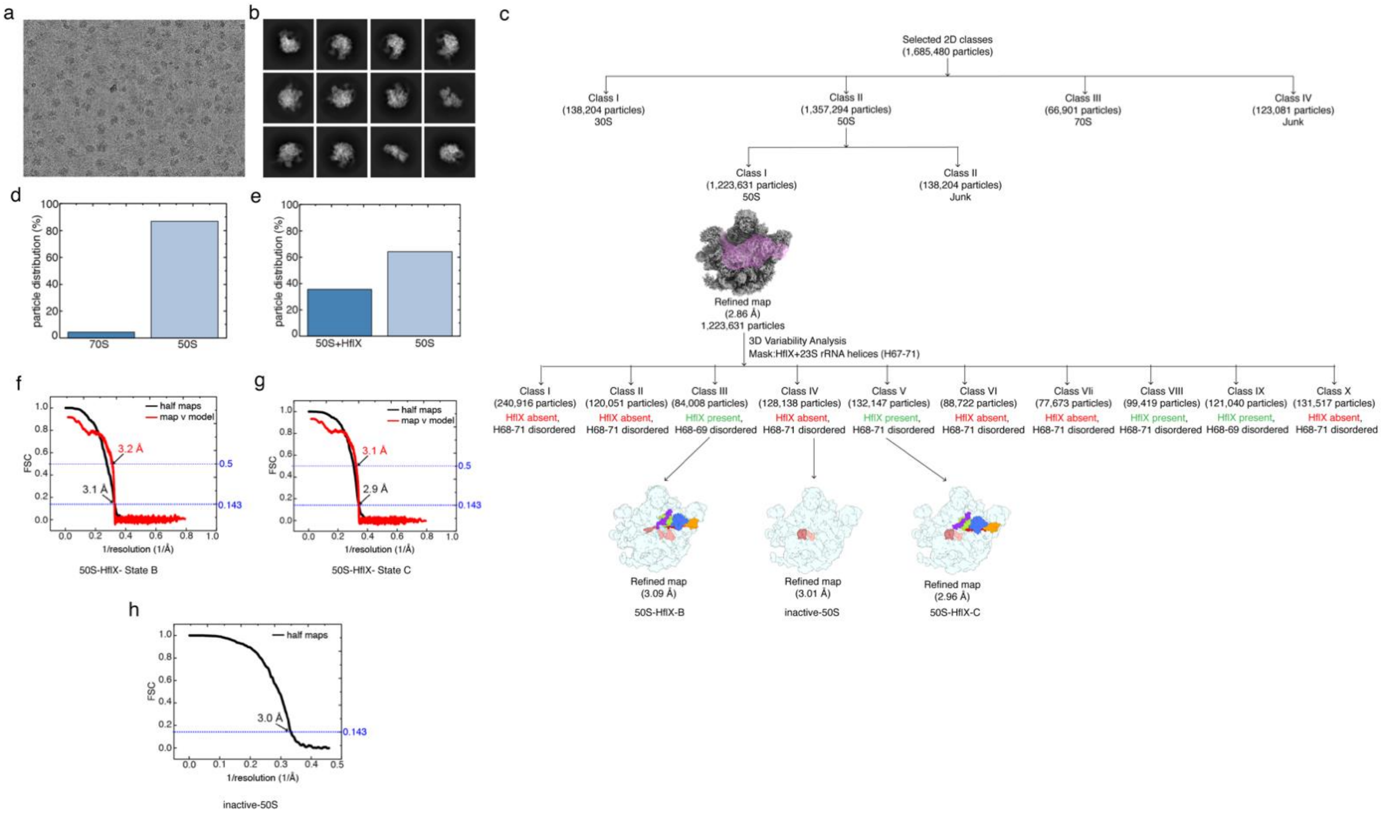


**Supplementary Figure 5.** Image processing of the *M. smegmatis* 70S-HflX complex prepared with higher concentration of HflX:GMPPNP. (**a**) a representative micrograph from the dataset. (**b**) Representative 2D-class averages selected for subsequent data-processing. (**c**) Flowchart showing the details of 3D classifications, 3D variability analysis and refinements performed on the selected particles. The final maps (50S-HflX-B, 50S-HflX-C and inactive 50S) used for model building and interpretations (in main Figures 2 and 3) are displayed. (**d**) Plot comparing the percentages of 70S and 50S ribosomal subunit obtained. (**e**) Plot comparing the percentages of HflX bound and unbound 50S ribosomal subunits. Gold-standard FSC (black) overlaid with the map-to-model FSC (red) for (**f**) the 50S-HflX- B and (**g**) the 50S-HflX-C. (**h**) Gold-standard FSC (black) plot for the inactive-50S map.

**Supplementary Figure 6.**

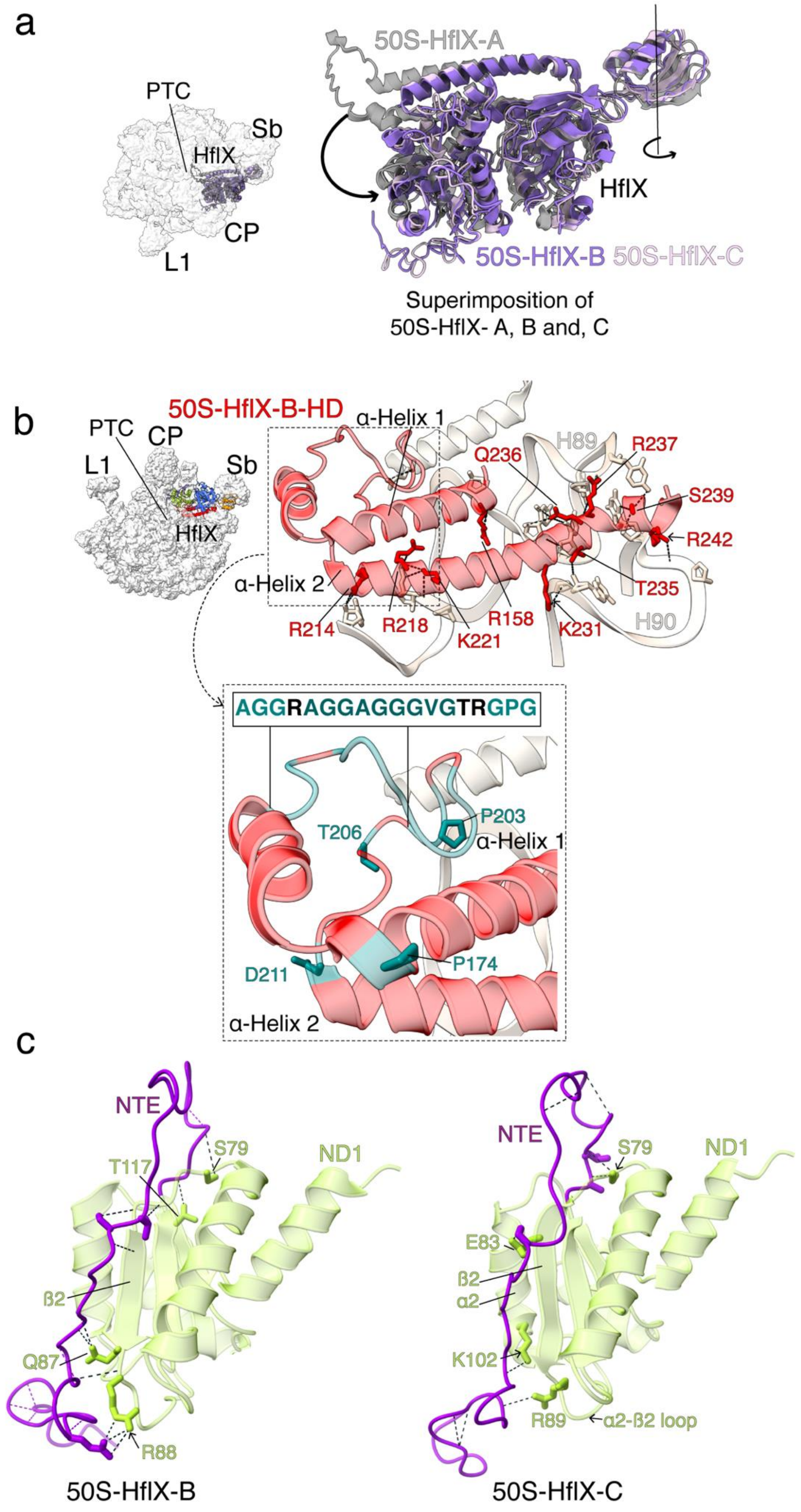

**Supplementary Figure 6.** (**a**) Superimposition of 50S-HflX -A, -B and -C shows that HflX in states B and C is deflected outwards by 3.4º, away from the 50S ribosomal subunit, about an axis passing through the CTD. (**b**) Molecular interactions of the retracted conformation of the HD in the 50S-HflX-B with the neighboring 23S rRNA helices, H89 and H90. The inset marks the point of origin of the bent conformation of helix I of HD at P174 (teal) that is followed by a stretch of glycines and alanines (sequence shown). The uncoiling in helix 2 of HD between T206 and R213 is also shown. (**c**) Molecular interactions of the HflX-NTE in 50S-HflX-B with ND1 (ß2 strand) and in C with ND1 (α2 helix, ß2 strand and the loop connecting α2-ß2) are displayed.

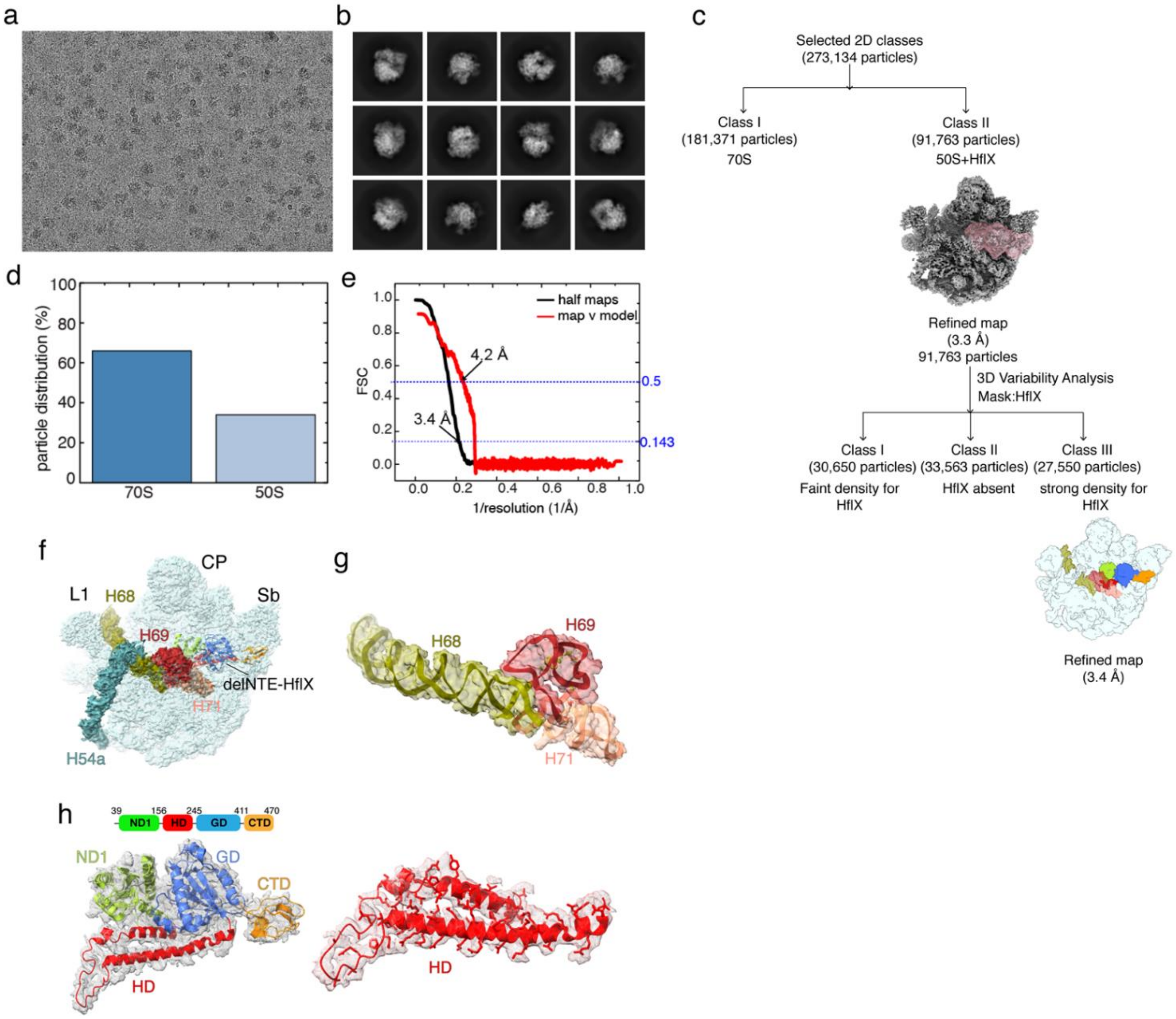


**Supplementary Figure 7.** Image processing of the *M. smegmatis* 70S-ΔNTE HflX complex and structure of the 50S-ΔNTE HflX complex. (**a**) a representative micrograph from the dataset. (**b**) Representative 2D-class averages selected for subsequent data-processing. (**c**) Flowchart showing the details of 3D classifications, 3D variability analysis and refinements performed on the selected particles. The final map of the 50S-ΔNTE HflX used for model building and interpretations is displayed. (**d**) Plot comparing the percentages of 70S monosome and 50S ribosomal subunit. (**e**) Gold-standard FSC (black) overlaid with the map-to-model FSC (red) plot for the 50S-ΔNTE HflX map. **f**) 3.4 Å cryo-EM structure of the 50S-ΔNTE HflX complex. Landmarks of the 50S ribosomal subunit, L1, L1 stalk; CP, central protuberance, and Sb, L7/L12 stalk base. The 23S rRNA helices H68 (olive green), H69 (brown) and H71 (light salmon) are highlighted. The HflX domains are color coded as in the main figure. (**g**) The well-resolved densities for the fully intact 23S rRNA helices H68, H69, and H71 can be seen. (**h**) Modeling of the ΔNTE-HflX into corresponding cryo-EM density. Model and density of the HflX-HD domain is shown (right) to emphasize that the HflX-HD is in an extended conformation.

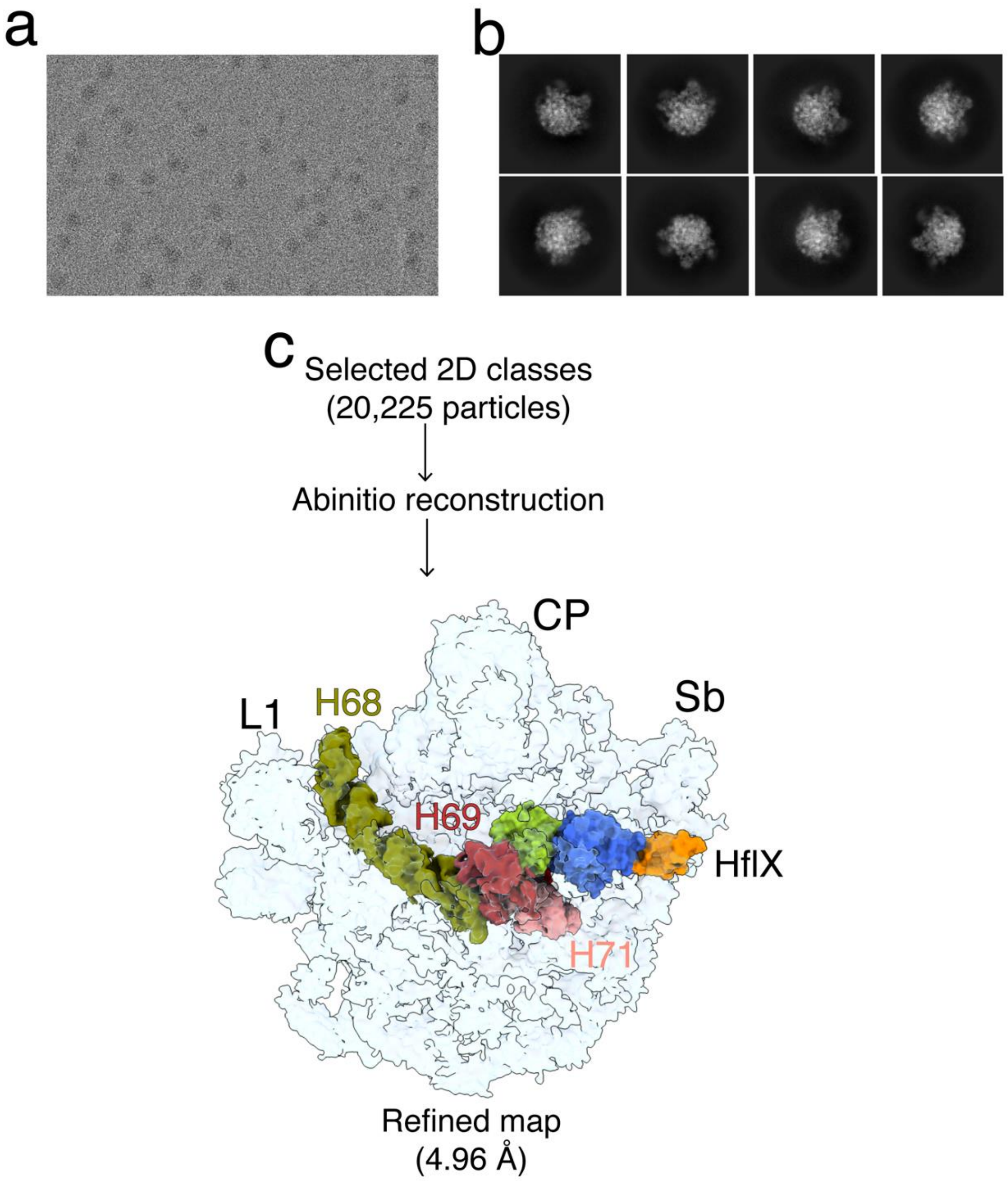


**Supplementary Figure 8.** Image processing of the pre-dissociated *M. smegmatis* 50S-HflX complex. (**a**) A representative micrograph from the dataset. (**b**) Representative 2D-class averages selected for subsequent data-processing. (**c**) Flowchart showing how the 50S-HflX map was reconstructed from the particles selected from 2D-classification. The final map with an overall resolution of 4.96 Å is displayed. Densities corresponding to H68 (olive green), H69 (brown), H71 (light salmon) and HflX (ND1, green; GD, blue; CTD, orange) are indicated.

### Supplementary Figure 9.

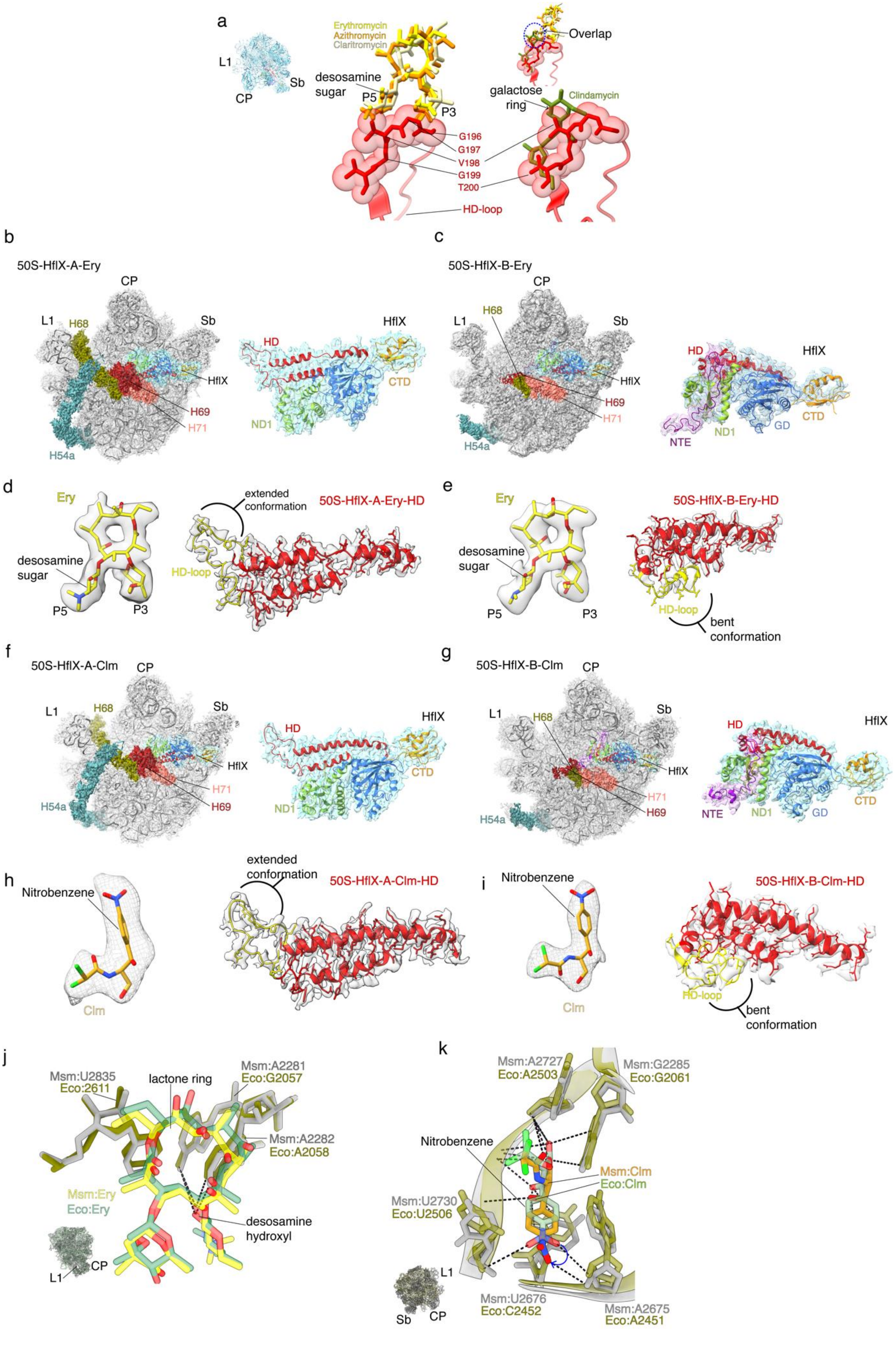

**Supplementary Figure 9. PTC drugs can bind to 50S in the presence of HflX by avoiding potential steric clash with HflX-HD.** (**a**) Superimposition of the *Msm* 50S-HflX-A structure with the structures of *Eco* 50S-erythromycin (yellow, PDB:3J5L), *Eco* 50S-azithromycin (orange, PDB:8E42) and *Mtu* 50S-clarithromycin (wheat, PDB: 7F0D) complexes reveals that the *Msm* HflX-HD loop would sterically clash with the sugar-moieties at positions 3 and 5 (P3, P5) of the antibiotics. Superimposition of the *Msm* 50S-HflX-A structure with that of the *Eco* 50S-clindamycin complex (olive green, PDB:8CGD) also shows a significant overlap with the *Msm* HflX-HD loop. **(b)** Cryo-EM structure of the *Msm* 50S-HflX-A-Ery complex. Landmarks of the 50S ribosomal subunit are as in main Figure 1 or Extended Figure 8. Additionally, the 23S rRNA helices H68 (olive green), H69 (brown), H71 (light salmon) and H54a (teal) are marked. Modeling of the HflX domain into their corresponding cryo-EM density. All the domains of HflX, except the NTE, are resolved. **(c)** Cryo-EM structure of the *Msm* 50S-HflX-B-Ery complex. Fitting of the HflX domain models into their corresponding cryo-EM density. All the domains of HflX, including the NTE, are resolved. **(d)** Densities corresponding to erythromycin (Ery) and the extended HflX HD-loop in the 50S-HflX-A-Ery structure, are displayed. The HD-loop is colored yellow while the HD-helices are colored in red. **(e)** Densities corresponding to Ery and the bent HflX HD-loop in the 50S-HflX-B-Ery structure are displayed. **(f and g)** Cryo-EM structure of the *Msm* 50S-HflX-A-Clm and 50S-HflX-B-Clm complexes. Modeling of the HflX domains into their corresponding cryo-EM densities. HflX-NTE is resolved in the 50S-HflX-B-Clm structure but not in 50S-HflX-A-Clm. (**h and i)** Densities corresponding to chloramphenicol (Clm) and the extended and bent HflX HD-loop, in the 50S-HflX-A-Clm and 50S-HflX-B-Clm structures, are displayed. **(j)** Superimposition of Msm 50S-HflX-B-Ery structure (gray) with that of Ec 50S-Ery (PDB:3J5L, in olive green) shows that Ery (yellow in Msm, green in Ec) is stabilized by similar interactions at the respective NPETs. **(k)** Superimposition of Msm 50S-HflX-B-Clm (gray) and Ec-70S-Clm (PDB:4V7T, in olive green) structures. Although Clm binding site is same in both structures, a rotation of the nitro group (blue arrow) of the nitrobenzene ring, along with a displacement of the nitrobenzene ring of the drug, are observed in the Msm structure (gold) as compared to Ec (green).

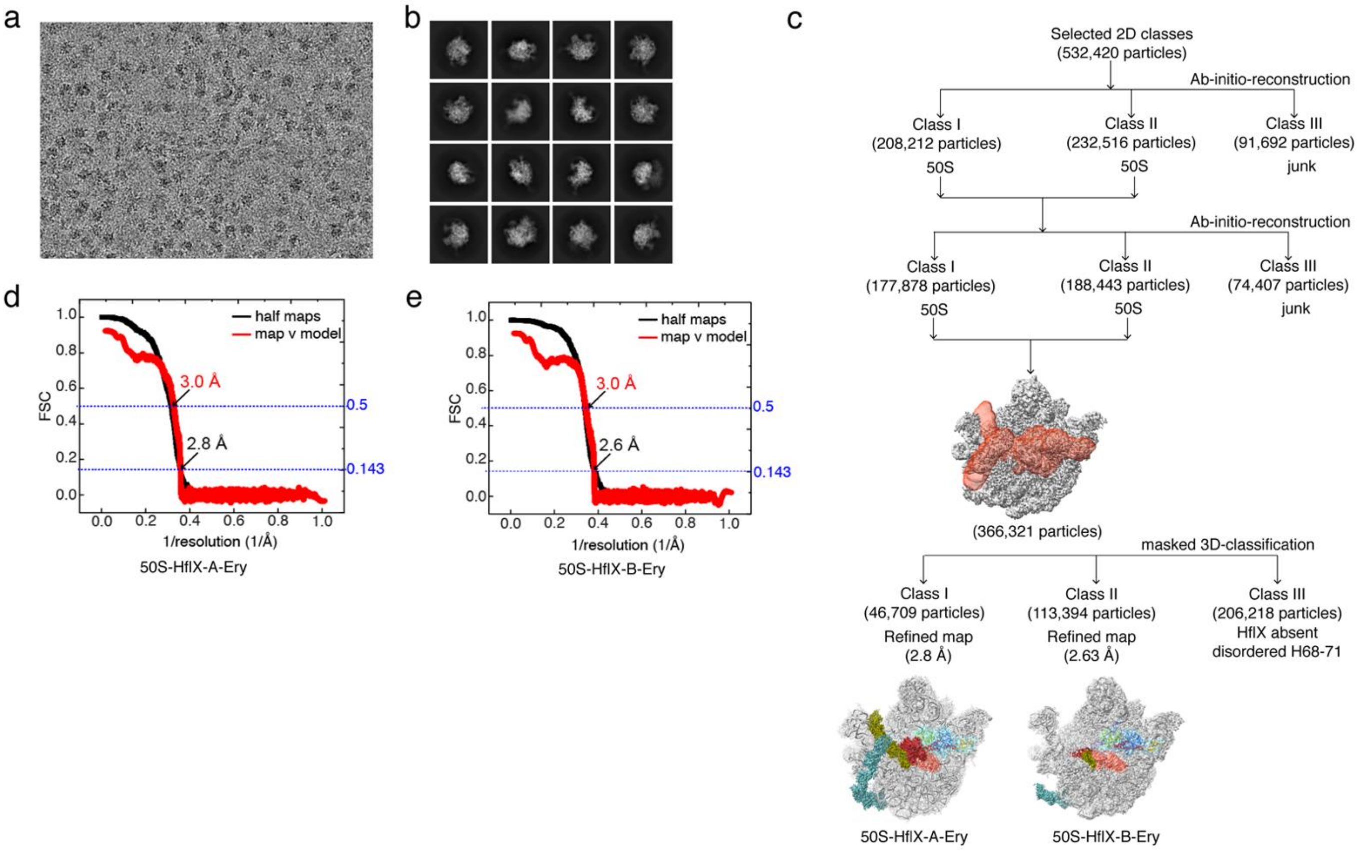


**Supplementary Figure 10.** Image processing of the Msm 70S-HflX-Ery complex. (**a**) A representative micrograph from the dataset. (**b**) A subset of the 2D-class averages selected for subsequent data-processing. (**c**) Flowchart showing how the 50S-HflX-A-Ery and 50S-HflX-B-Ery maps were reconstructed from the particles selected from 2D-classification. The final maps with an overall resolution of 2.8 Å and 2.63 Å are displayed. Densities corresponding to H68 (olive green), H69 (brown), H71 (light salmon) and HflX (ND1, green; GD, blue; CTD, orange) are indicated.

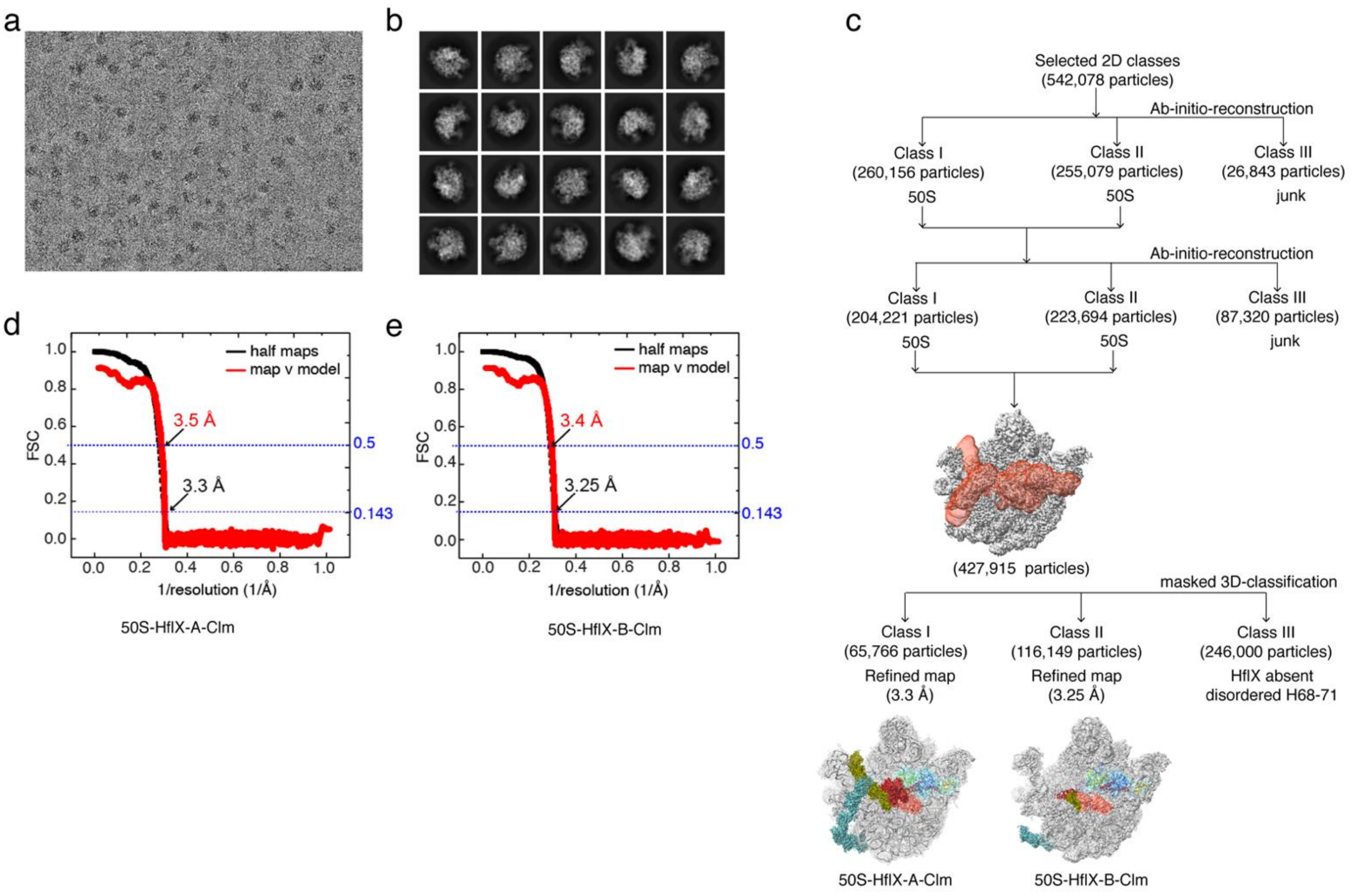


**Supplementary Figure 11.** Image processing of the Msm 70S-HflX-Clm complex. (**a**) A representative micrograph from the dataset. (**b**) A subset of the 2D-class averages selected for subsequent data-processing. (**c**) Flowchart showing how the 50S-HflX-A-Clm and 50S-HflX-B-Clm maps were reconstructed from the particles selected from 2D-classification. The final maps with an overall resolution of 3.3 Å and 3.25 Å are displayed. Densities corresponding to H68 (olive green), H69 (brown), H71 (light salmon) and HflX (ND1, green; GD, blue; CTD, orange) are indicated.

**Supplementary Figure 12.**

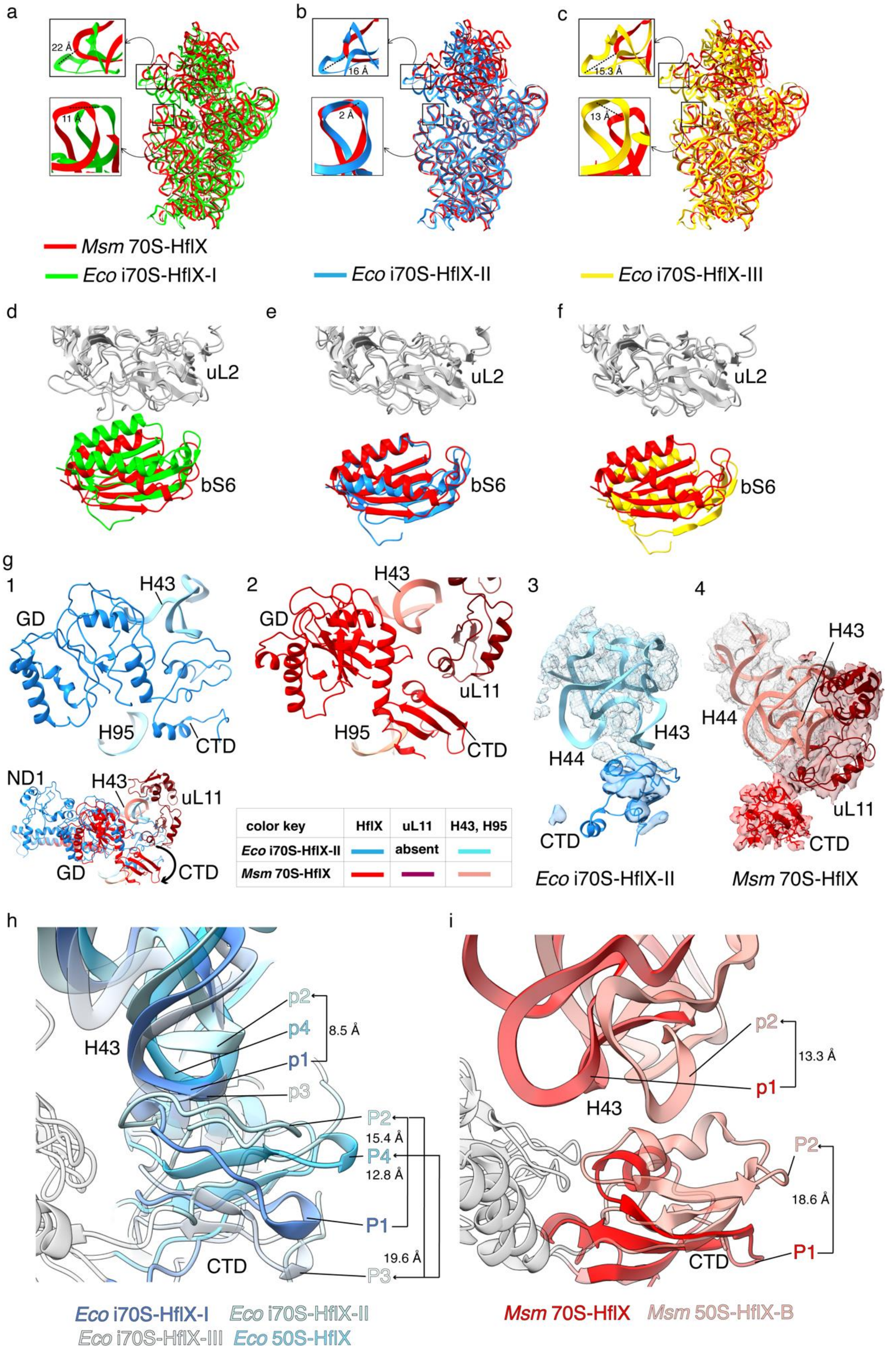

**Supplementary Figure 12**. **Comparison between structures of *Msm* 70S-HflX (red) and *Eco* 70S-HflX intermediates.** (a) *Eco*-i70S-HflX-I (green, PDB: 8G34), (b) *Eco*-i70S-HflX-II (deep sky blue, PDB: 8G31), and (c) *Eco*-i70S-HflX-III (yellow, PDB: 8G38). The insets illustrate the relative displacement of the rRNA at the SSU beak and shoulder. Comparison of the relative displacement of the SSU protein bS6 relative to LSU protein uL2 between *Msm* 70S-HflX (red) and the *Eco* 70S-HflX intermediates: (d) *Eco*-i70S-HflX-I (green, PDB: 8G34), (e) *Eco*-i70S-HflX-II (deep sky blue, PDB: 8G31), and (f) *Eco*-i70S-HflX-III (yellow, PDB: 8G38). (g) The thumbnail depicts the superimposition of *Msm* 70S-HflX and Eco-i70S-HflX-II to illustrate the differences in HflX domains and the Sb components (H43 and uL11). (g1) The positions of the HflX-CTD and HflX-GD with respect to H43 and H95 in *Eco*, (g2) the position of the HflX-CTD and HflX-GD with respect to H43, H95, and uL11 in *Msm*, (g3) cryo-EM density corresponding to HflX-CTD and the Sb in *Eco*-i70S-HflX-II (EMD-29687), and (g4) *Msm*. (h) Superimposition of the three TRCEM-derived *Eco*-70S-HflX intermediates and *Eco*-50S-HflX (PDB: 5ADY) to illustrate the relative motion of HflX-CTD with respect to H43 (in shades of blue). The different positions of HflX-CTD (uppercase 'P') and H43 (lowercase 'p') are marked, and their displacements from one position to the next are indicated. (i) Superimposition of *Msm*-70S-HflX and *Msm*-50S-HflX-B structures to illustrate the relative motion of HflX-CTD with respect to H43 (in red and salmon, respectively). The two positions of HflX-CTD (uppercase 'P') and H43 (lowercase 'p') are marked, and their displacements from their initial to final positions are indicated by arrows.

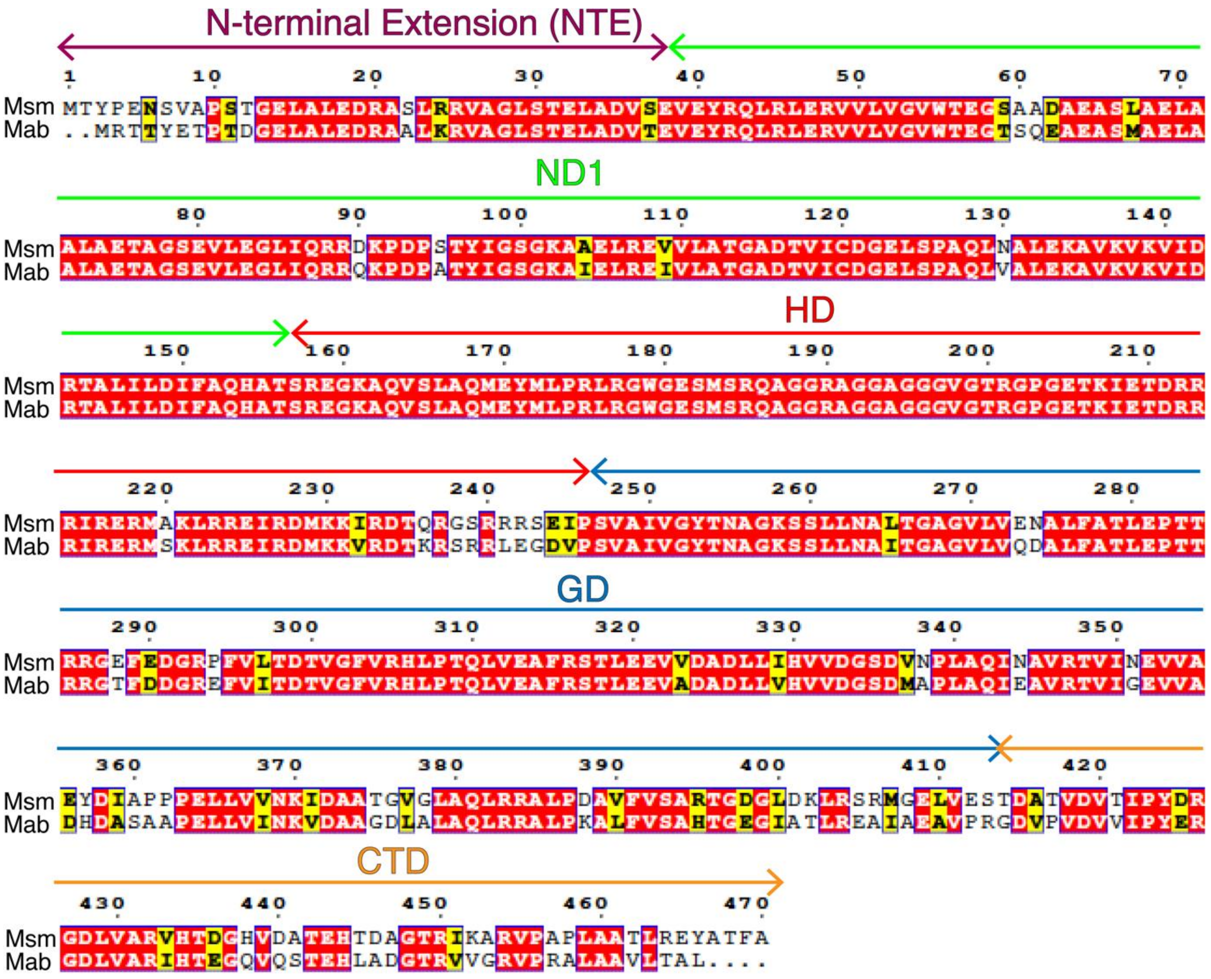


**Supplementary Figure 13.** Sequence alignment of HflX homologs from *Mycobacterium smegmatis* (Msm) and *Mycobacterium abscessus* (Mab). Identical (red) and similar (yellow) regions are highlighted, and domain boundaries are identified.

**Supplementary Table 1. Grid preparation, data collection and refinement parameters for the ribosome-HflX complexes.**

| | **70S-HflX** | **50S-HflX-A** | **50S-HflX-B** | **50S-HflX-C** | **50S-ΔNTE-HflX** |
|---|---|---|---|---|---|
| PDB | 8VIO | 8VK0 | 8VK7 | 8VKI | 8VKW |
| EMBD | EMD-43267 | EMD-43294 | EMD-43305 | EMD-43317 | EMD-43333 |
| Grid | | | | | |
| Hole size/spacing | R1.2/1.3 | R1.2/1.3 | R1.2/1.3 | R1.2/1.3 | R1.2/1.3 |
| Mesh | 300 (Cu) | 300 (Cu) | 300 (Cu) | 300 (Cu) | 300 (Cu) |
| Volume (µl) | 3 | 3 | 3 | 3 | 3 |
| Blot time (s) | 5 | 5 | 5 | 5 | 5 |
| Temperature (°C) | 4 | 4 | 4 | 4 | 4 |
| Humidity (%) | 100 | 100 | 100 | 100 | 100 |
| **Data Collection** | | | | | |
| Microscope | FEI Titan Krios | FEI Titan Krios | FEI Titan Krios | FEI Titan Krios | FEI Titan Krios |
| Electron source | Field Emission Gun | Field Emission Gun | Field Emission Gun | Field Emission Gun | Field Emission Gun |
| Acceleration voltage (keV) | 300 | 300 | 300 | 300 | 300 |
| Camera | GatanK3 | GatanK3 | GatanK3 | GatanK3 | GatanK3 |
| Nominal magnification | 81,000 | 81,000 | 64,000 | 64,000 | 81,000 |
| Pixel size | 1.0825 | 1.0825 | 1.076 | 1.076 | 0.846 |
| Total electron exposure ($e^-/Å^2$) | 51.22 | 51.22 | 51.85 | 51.85 | 60.64 |
| Nominal minimum defocus (nm) | 1800 | 1800 | 1000 | 1000 | 800 |
| Nominal maximum defocus (nm) | 2500 | 2500 | 2500 | 2500 | 2500 |
| Number of collected micrographs | 15,011 | 15,011 | 8,934 | 8,934 | 8,700 |
| Number of extracted particles | 1,143,540 | 1,143,540 | 2,585,153 | 2,585,153 | 332,311 |
| Number of complex particles | 48,443 | 57,434 | 84,008 | 132,147 | 27,550 |
| Resolution (Å) | 3.26 | 3.14 | 3.09 | 2.96 | 3.44 |
| FSC threshold | 0.143 | 0.143 | 0.143 | 0.143 | 0.143 |
| Model-to-map | | | | | |
| CC mask | 0.62 | 0.62 | 0.66 | 0.60 | 0.71 |
| CC volume | 0.63 | 0.63 | 0.67 | 0.61 | 0.71 |
| Resolution (Å) | 3.5 | 3.3 | 3.3 | 3.1 | 4.2 |
| FSC threshold | 0.5 | 0.5 | 0.5 | 0.5 | 0.5 |
| **Model** | | | | | |
| Atoms | 152,629 | 100,929 | 96,832 | 96,565 | 97,622 |
| Nucleotide residues | 4,825 | 3,237 | 4,050 | 3,070 | 3,161 |
| Protein residues | 6,334 | 4,107 | 3,064 | 4,001 | 3,883 |
| Bonds (RMSD) | | | | | |
| Length (Å) | 0.006 | 0.009 | 0.006 | 0.008 | 0.005 |
| Angles (°) | 0.795 | 1.099 | 0.657 | 0.767 | 0.607 |
| MolProbity score | 2.08 | 2.16 | 2.19 | 2.15 | 2.13 |
| Clash score | 11.95 | 7.77 | 12.97 | 12.79 | 12.3 |
| **Ramachandran plot** | | | | | |
| Outliers | 0.06 | 0.05 | 0.00 | 0.05 | 0.03 |
| Allowed | 8.02 | 10.97 | 10.62 | 9.33 | 9.33 |
| Favored | 91.92 | 88.98 | 89.38 | 90.62 | 90.65 |
| Ramachandran plot Z-score, whole | -2.52 | -4.57 | -2.40 | -2.10 | -2.26 |
| Rotamer outliers (%) | 0.89 | 1.62 | 0.56 | 0.79 | 0.00 |
| Cß outliers (%) | 0.02 | 0.03 | 0.00 | 0.06 | 0.00 |
| CαBLAM outliers (%) | 3.93 | 5.87 | 5.29 | 4.68 | 4.53 |
| **ADP B factors, mean ($Å^2$)** | | | | | |
| Protein | 35.53 | 75.72 | 68.51 | 59.47 | 276.9 |
| Nucleotide | 47.08 | 63.94 | 73.67 | 76.71 | 249.07 |

### Supplementary Table 2. Grid preparation, data collection and refinement parameters*

| | 70S-P-tRNA | Inactive-50S | Pre-dissociated 50S-HflX |
|---|---|---|---|
| EMBD | EMD-43791 | EMD-43778 | EMD-44044 |
| Grid | | | |
| Hole size/spacing | R1.2/1.3 | R1.2/1.3 | R1.2/1.3 |
| Mesh | 300 (Cu) | 300 (Cu) | 300 (Cu) |
| Volume (µl) | 3 | 3 | 3 |
| Blot time (s) | 5 | 5 | 5 |
| Temperature (ºC) | 4 | 4 | 4 |
| Humidity (%) | 100 | 100 | 100 |
| **Data Collection** | | | |
| Microscope | FEI Titan Krios | FEI Titan Krios | JEOL 3200FSC |
| Electron source | Field Emission Gun | Field Emission Gun | Field Emission Gun |
| Acceleration voltage (keV) | 300 | 300 | 300 |
| Camera | GatanK3 | GatanK3 | GatanK2 Summit |
| Nominal magnification | 81,000 | 64,000 | 25,000 |
| Pixel size | 1.0825 | 1.076 | 1.25 |
| Total electron exposure ($e^-/Å^2$) | 51.22 | 51.85 | 21 |
| Nominal minimum defocus (nm) | 1800 | 1000 | 500 |
| Nominal maximum defocus (nm) | 2500 | 2500 | 3500 |
| Number of collected micrographs | 15,011 | 8,934 | 831 |
| Number of extracted particles | 1,143,540 | 2,585,153 | 168,372 |
| Number of complex particles | 188,044 | 128,138 | 20,225 |
| Resolution (Å) | 3.06 | 3.0 | 4.96 |
| FSC threshold | 0.143 | 0.143 | 0.143 |

**These three reconstructions were obtained for qualitative comparison as control, and atomic models for these maps were not submitted.*

**Supplementary Table 3. Grid preparation, data collection and refinement parameters for antibiotic-bound ribosome-HflX complexes.**

| | **50S-HflX-A-Ery** | **50S-HflX-B-Ery** | **50S-HflX-A-Clm** | **50S-HflX-B-Clm** |
|---|---|---|---|---|
| PDB | 8VR4 | 8VPK | 8VRL | 8VR8 |
| EMBD | EMD-43476 | EMD-43409 | EMD-43484 | EMD-43477 |
| Grid | | | | |
| Hole size/spacing | R1.2/1.3 | R1.2/1.3 | R1.2/1.3 | R1.2/1.3 |
| Mesh | 300 (Cu) | 300 (Cu) | 300 (Cu) | 300 (Cu) |
| Volume (µl) | 3 | 3 | 3 | 3 |
| Blot time (s) | 5 | 5 | 5 | 5 |
| Temperature (°C) | 4 | 4 | 4 | 4 |
| Humidity (%) | 100 | 100 | 100 | 100 |
| **Data Collection** | | | | |
| Microscope | FEI Titan Krios | FEI Titan Krios | FEI Titan Krios | FEI Titan Krios |
| Electron source | Field Emission Gun | Field Emission Gun | Field Emission Gun | Field Emission Gun |
| Acceleration voltage (keV) | 300 | 300 | 300 | 300 |
| Camera | GatanK3 | GatanK3 | GatanK3 | GatanK3 |
| Nominal magnification | 81,000 | 81,000 | 105,000 | 105,000 |
| Pixel size | 0.846 | 0.846 | 0.8443 | 0.8443 |
| Total electron exposure ($e^-$/Å$^2$) | 62.87 | 62.87 | 70.14 | 70.14 |
| Nominal minimum defocus (nm) | 1000 | 1000 | 800 | 800 |
| Nominal maximum defocus (nm) | 1600 | 1600 | 2500 | 2500 |
| Number of collected micrographs | 12,145 | 12,145 | 14,374 | 14,374 |
| Number of extracted particles | 1,747,436 | 1,747,436 | 2,117,095 | 2,117,095 |
| Number of complex particles | 46,709 | 113,394 | 65,766 | 116,149 |
| Resolution (Å) | 2.8 | 2.63 | 3.33 | 3.25 |
| FSC threshold | 0.143 | 0.143 | 0.143 | 0.143 |
| Model-to-map | | | | |
| CC mask | 0.55 | 0.45 | 0.57 | 0.52 |
| CC volume | 0.52 | 0.46 | 0.58 | 0.54 |
| Resolution (Å) | 3.0 | 3.0 | 3.5 | 3.4 |
| FSC threshold | 0.5 | 0.5 | 0.5 | 0.5 |
| **Model** | | | | |
| Atoms | 99,355 | 96,925 | 96,922 | 93,099 |
| Nucleotide residues | 3,236 | 4,053 | 3,237 | 3,068 |
| Protein residues | 3,890 | 3,066 | 3,575 | 3,551 |
| Bonds (RMSD) | | | | |
| Length (Å) | 0.006 | 0.004 | 0.006 | 0.003 |
| Angles (°) | 0.713 | 0.736 | 0.760 | 0.677 |
| MolProbity score | 2.16 | 2.10 | 1.95 | 1.75 |
| Clash score | 14.16 | 14.23 | 7.57 | 5.37 |
| **Ramachandran plot** | | | | |
| Outliers | 0.08 | 0.08 | 0.06 | 0.00 |
| Allowed | 8.45 | 6.80 | 9.30 | 7.42 |
| Favored | 91.47 | 93.12 | 90.65 | 92.58 |
| Ramachandran plot Z-score, whole | -1.73 | -0.90 | -2.77 | -1.95 |
| Rotamer outliers (%) | 0.13 | 0.06 | 0.56 | 0.32 |
| Cß outliers (%) | 0.03 | 0.03 | 0.00 | 0.00 |
| CαBLAM outliers (%) | 3.73 | 3.55 | 5.14 | 4.81 |
| **ADP B factors, mean (Å$^2$)** | | | | |
| Protein | 34.33 | 18.23 | 24.06 | 25.02 |
| Nucleotide | 48.26 | 27.21 | 44.00 | 39.38 |
